# Highly structured, partner-sex- and subject-sex-dependent cortical responses during social facial touch

**DOI:** 10.1101/545434

**Authors:** Christian L. Ebbesen, Evgeny Bobrov, Rajnish P. Rao, Michael Brecht

**Affiliations:** Bernstein Center for Computational Neuroscience Berlin, Humboldt-Universität zu Berlin, 10115 Berlin, Germany; Berlin School of Mind and Brain, Humboldt-Universität zu Berlin, 10115 Berlin, Germany; Neuroscience Institute, New York University, NY 10016, USA; QUEST Center for Transforming Biomedical Research, Berlin Institute of Health, 10178 Berlin, Germany; NeuroCure Cluster of Excellence, Humboldt-Universität zu Berlin, 10115 Berlin, Germany

## Abstract

Touch is a fundamental aspect of social, parental and sexual behavior. Despite our detailed knowledge about cortical processing of non-social touch, we still know little about how social touch cortical circuits. We investigated neural activity across five frontal, motor and sensory cortical areas in rats engaging in naturalistic social facial touch. Information about social touch and the sex of the interaction partner (a biologically significant feature) is a major determinant of cortical activity. 25.3% of units were modulated during social touch and 8.3% of units displayed ‘sex-touch’ responses (responded differently, depending on the sex of the interaction partner). Single-unit responses were part of a structured, partner-sex- and, in some cases, subject-sex-dependent population response. Simulations suggest that a change in inhibitory drive might underlie these population dynamics. Our observations suggest that socio-sexual characteristics of touch (subject and partner sex) widely modulate cortical activity and need to be investigated with cellular resolution.

## Introduction

Social touch is a powerful emotional stimulus^1–4^. Harnessing the ability of social touch to modulate emotion for example by caress or massage has emerged as a protective and therapeutic strategy for various mental health conditions, such as anxiety, depression and autism spectrum disorder^3–7^. However, despite our detailed knowledge about cortical processing of discriminative, non-social touch, we still know very little about how social touch modulates cortical circuits.

Human social touch is different from non-social touch, in part due to activation of a specialized ‘bottom-up’ pathway. Light touch and caress activates a particular class of C-tactile mechanosensory afferents in hairy skin^3,8^. Activation of these fibers leads to decreased levels of the stress hormone cortisol, mediates release of the neuropeptide oxytocin and modulates the activity of insular cortex, a key region for emotional regulation^9–12^. However, two major lines of evidence also point to a major role of ‘top-down’ modulation in the processing of social touch.

First, unlike discriminative touch, the psychophysics of social touch is extremely dependent on individual, social and sexual context. For example, even though the actual tactile input is essentially identical, the touch of a loved one can feel comforting and pleasant while the touch of someone repulsive can feel intensely aversive^4,8,13^. Second, activation of the C-tactile fibers is not nescessary to make social touch ‘special’. Interpersonal touch delivered by the glabrous skin of the palm, thus not activating the C-tactile fibers, elicits a social softness illusion in the giver^14^ and has a different neural activation pattern from non-social touch^15–17^.

Pioneering functional imaging studies in humans have investigated which cortical regions contribute to the top-down modulation of social touch processing by identifying brain regions, where activity patterns depend on the social ‘meaning’ of the touch, rather than the actual haptic input. For example, although in fact an identical pattern of touch was always given by the same experimenter, activity in anterior cingulate, orbitofrontal, somatosensory and insular cortices is different when subjects believe they are being touched by a man or a woman^18,19^ or by a partner or a stranger^20^. This social-context-dependent modulation of cortical touch responses is negatively correlated with autism-like traits^19^ and is increased by intranasal oxytocin administration^20^. In line with a role of these regions in top-down modulation of social touch processing, somatosensory cortex, for example, is activated in a social context before any actual touch input^16,17^, when observing others being touched ^21^, and when simply imagining pleasant or sexual touch^22^.

In this paper, we apply techniques with cellular resolution in the rat, which has been pioneered in the primate by Romo and colleagues^23–26^, namely to investigate and compare how single cortical neurons across multiple cortical areas respond during the same somatosensory stimulus. We focus on rat social facial touch, during which rats align their snouts and palpate each other’s faces with their whiskers^27^. Social facial touch has attractive behavioral characteristics. First, the behavior is ecologically valid. The animals are untrained, social interactions are jointly initiated by the animals themselves and the animals are freely moving, thus their cortical activity presumably closely resembles activity in a natural social setting. Second, by letting animals interact with partners of both sexes, we can manipulate the social context of the touch, while keeping the actual haptic input identical or similar. The socio-sexual ‘meaning’ of touching male and female conspecifics is different^28^, but during social facial interactions, male rats whisk with equal power onto conspecifics of both sexes. Females whisk onto females like a male rat would, but whisk with lower whisking amplitudes onto males^27^.

Similar to humans, previous work has shown that even though whisking amplitude is lower during social facial interactions than when investigating objects, population firing rate changes^29^ and membrane potential modulations^30^ in rat somatosensory cortex are larger during social touch than object touch and do not correlate with whisker movements. Also similar to humans, rat somatosensory activity is modulated in a social context before actual social facial touch^30^ and firing rates depends on socio-sexual context, such as estrus state^29,31^ and emotional state^32^.

We present a flexible regression approach, which allows us to ask how social touch impacts activity, despite the large behavioral variability in natural social interactions. We ask the following questions: (i) How is information about touch and social context (partner sex) represented at the level of single neurons? (ii) How widely is this information available across five different cortical areas? (iii) How does social context impact population dynamics during touch? (iv) How does the population response structure depend on cortical area, partner sex and sex of the subject animal itself? (v) Which cellular and network mechanisms could plausibly underlie cortical population dynamics during social touch?

We present a flexible regression approach that allowed us to investigate how social context (partner sex and subject sex) impacts cortical processing during naturalistic social touch, an untrained, self-initiated social behavior with no imposed trial structure. Social context (partner sex) modulates the firing patterns of individual cortical neurons, and information about social touch and partner sex is available across cortical areas with diverse functions (somatosensory, auditory, motor, frontal) and diverse response patterns (mostly increasing or mostly decreasing during touch). Firing rate changes of single neurons are not random across the network. On the contrary, cortical networks respond to social touch with highly structured, partner-sex and – in some cases – subject-sex-dependent population responses. Network simulations of cortical touch responses reveal that a change in inhibitory drive might underlie these population dynamics.

## Results

### Data

In this study, we analyze 1,156 single neurons, recorded from five cortical areas, over 7,408 episodes of social facial touch in 15 female and 14 male rats (58,591 unique cell-touch pairs, averaging 51 touch episodes per cell).

### Behavioral variability and diverse cortical responses during social facial touch

To investigate how social touch modulates cortical processing across frontal and sensory cortices, we implanted tetrodes to record single-unit responses from freely moving, socially interacting male and female rats in the ‘social gap paradigm’ (Figure 1a). In this paradigm, rats reach across a gap between elevated platforms to engage in social facial touch; an un-trained, self-initiated social behavior where the animals align their snouts and palpate each other’s faces with their whiskers^27^.

We recorded the activity of neurons throughout the cortical column in five cortical areas: Two sensory areas, barrel cortex (‘SI’, the primary somatosensory representation of the mystacial vibrissae) and auditory cortex (‘Al’), and three frontal areas, vibrissa motor cortex (‘VMC’, the primary motor representation of the mystacial vibrissae), cingulate cortex (‘ACC’, a putative homolog of human anterior cingulate cortex) and prelimbic cortex (‘PrL’, a putative homolog of human medial prefrontal cortex) (Figure 1b). To investigate how cortical processing of social facial touch depends on the subject sex and the sex of the social interacting partner, we recorded from both male and female rats, interacting with multiple male and female conspecifics (see Methods). A portion of the data analyzed here has been presented in previous studies where we investigated other questions (ref.^29,33,34^, see Methods).

We plotted the activity of single neurons, across all social interactions, aligned to the onset of whisker-to-whisker touch (peri-stimulus time histograms, ‘PSTHs’). From inspecting these plots, we made two preliminary observations. First, we noticed that cortical responses to social facial touch are widespread and diverse: In all brain areas, we both found neurons, which significantly increased (Figure 1c) and significantly decreased (Figure 1d) their activity at the onset of social facial touch. Second, we noticed that the PSTHs were highly variable from trial to trial. This variability could reflect a low correlation between firing patters and behavior. On the other hand, if neural responses are tightly locked to behavior, but the behavior itself is highly variable, we would expect the same pattern. To discern these possibilities, we investigated the temporal statistics of the social facial touch episodes.

The median duration of a social facial touch was 1.33 s (IQR: 0.77-2.53 s, Figure 1e, top) and the median inter-touch-interval (ITI) was 4.68 s (IQR: 1.24-23.78 s, Figure 1e, middle), and both were highly variable. The duration of touch episodes varied across two orders of magnitude, the ITI varied across four orders of magnitude and the distributions of touch durations and inter-touch-intervals were highly overlapping (Figure 1e, bottom). In other words, social facial touch episodes happen in bouts where the animals engage in several short touches, separated by inter-touch-intervals, which are often on the same order of magnitude of the touch durations themselves. This behavioral observation suggests that a PSTH-based analysis (Figure 1c-d) will under-estimate the magnitude and overestimate the trial-to-trial varia-bility of neuronal responses to social facial touch. The ‘baseline’ period (−2000 to 0 ms) is not ‘clean’, so to speak, but con-taminated with touch episodes, and the post-stimulus period (0 to 500 ms) contains a mixture of trials with a wide range of touch durations.

**Figure 1.**
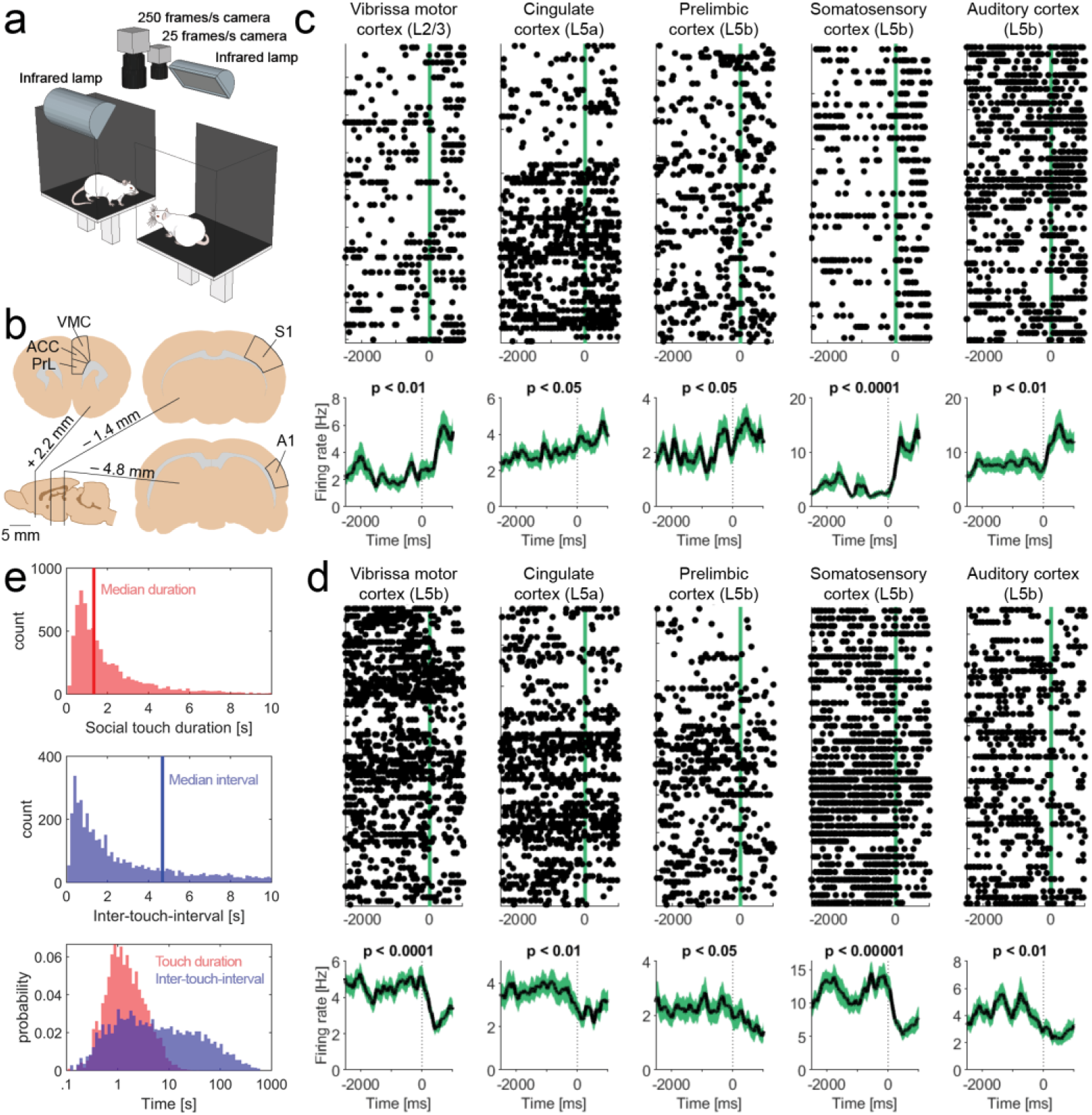
Diverse cortical responses and variable behavioral timing during social facial touch. **(a)** The social gap paradigm^27^. Rats separated on two platforms will reach across the gap to engage in social facial touch. In social facial touch, rats align their snouts and whisk to palpate each other’s face with their mystacial vibrissae. We recorded the behavior of male and female rats by videography from above in visual darkness under infrared illumination^29,33,34^. (b) Anatomical location of vibrissa motor cortex (‘VMC’ – the primary motor representation of the mystacial vibrissae), cingulate cortex (‘ACC’ – a putative homolog of human/primate anterior cingulate cortex), prelimbic cortex (‘PrL’ – a putative homolog of human/primate medial prefrontal cortex), barrel cortex (‘’ – the primary somatosensory representation of the mystacial vibrissae) and auditory cortex (‘A1’). (c) Example peri-stimulus time histograms (PSTHs) of touch-activated single neurons (Wilcoxon signed-rank test), aligned to the first whisker-to-whisker touch in each social touch episode. Raster plot above shows single trials, black dots indicate single spikes and vertical line indicates beginning of social facial touch. Black line below indicates mean firing rate, smoothed with an Alpha kernel (*τ* = 75 ms), shaded area indicates s.e.m. (d) Example PSTHs of touch-suppressed single neurons (Wilcoxon signed-rank test), aligned to the first whisker-to-whisker touch in each social touch episode. Same plotting convention as in (c). (e) Top: Histograms of touch duration (red) and inter-touch-time (blue) of social facial touch episodes. The time axes of both plots are clipped at 10 s. Below: Probability distributions of touch duration (red) and inter-touch-interval (blue) of social touch episodes, plotted on a logarithmic time scale. The two distributions span multiple orders of magnitude and strongly overlap.

### Single cortical neurons signal social touch and partner sex

To overcome the challenges presented by the highly variable temporal statistics of naturalistic social interactions, we used a generalized linear regression approach. Briefly, we modeled the spiking activity of the neurons as a Poisson process with history dependence and baseline fluctuations^35,36^ (graphical depiction of the data analysis and statistical modeling ‘pipe-line’ shown in Figure 2). We used a maximum likelihood approach combined with a non-parametric shuffling procedure to investigate the effect of social touch and the sex of the social interaction partner, while maintaining the information about duration and inter touch interval of every single touch episode (see Methods). We quantified the modulation of firing rates as a regression coefficient (a *β*-coeficient), which can be thought of as a modulation index that measures firing rate changes ‘fold changes’ (*β* = 1 corresponds to an e-fold increase, *β* = −1 corresponds to an e-fold decrease; motivation and an in-depth introduction to the use of beta-coefficients as a measure of firing rate changes across neural populations is provided in Suppl. Note 1). This approach allowed us to ask how social touch impacts neural activity, despite the behavioral variability.

**Figure 2.**
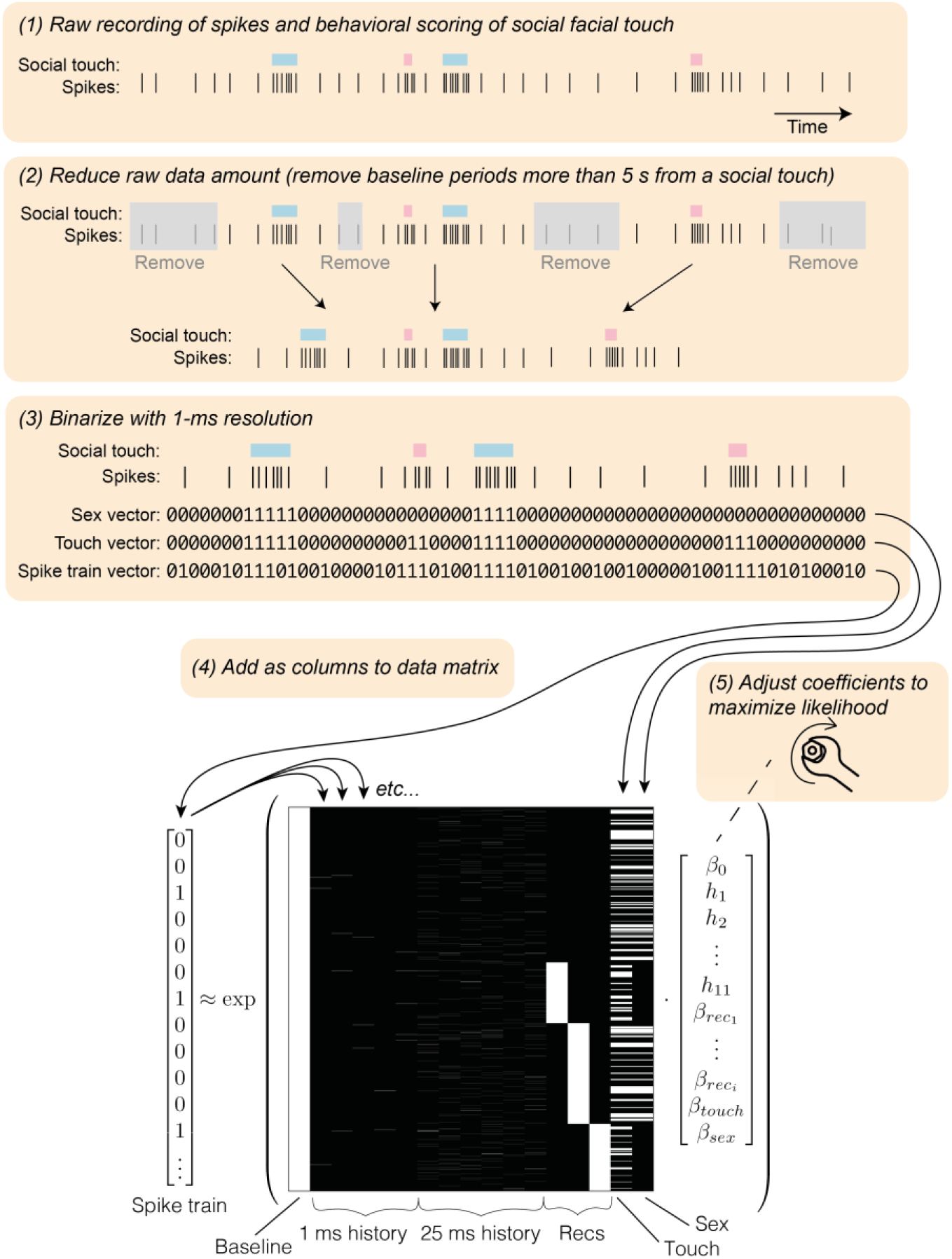
Graphical depiction of the spike train analysis. *Above:* Schematic depicting the five steps of the data analysis pipeline (cartoon data, with enlarged bin-size for visibility; real data is binned in 1-ms bins). *Below:* Example predictor matrix (real data) for the generalized linear regression approach. We discretize the spike train in 1-ms bins, reduce the amount of raw data (by removing baseline periods more than five seconds from a social interaction) and model the firing rate as a Poisson process^35,36^ (see Methods). The predictor matrix has the pressed touch neuron (a layer 5b neuron in following columns (left to right): a constant baseline rate; five 1 -ms spike history bins; six 25-ms history bins; three one-hot columns to model possible changes in baseline between the (in this example case) four recordings; a one-hot column indicating all social touch episodes; a one-hot column indicating the sex of the stimulus animal (0/1 corresponds to male/female, in this example case, the interaction partner animals were male in recording one and recording three). The vector indicates regression coefficients, which we fit numerically by likelihood maximization.

Using this statistical approach, we identified neurons that were modulated by social touch (‘touch neurons’), and neurons that were modulated by social touch, but responded differently when touching male and female conspecifics (‘sex-touch neurons’). Touch and sex-touch neurons had very diverse response patterns. We both found touch and sex-touch neurons that increased and decreased their firing rates during social touch (Figure 3a-e, left), and sex-touch neurons that responded more strongly to either female or male conspecifics (Figure 3a-e, right, additional examples of the diversity of sex-touch responses across the cortical areas are plotted in Supplementary Figure 1a-e).

**Figure 3.**
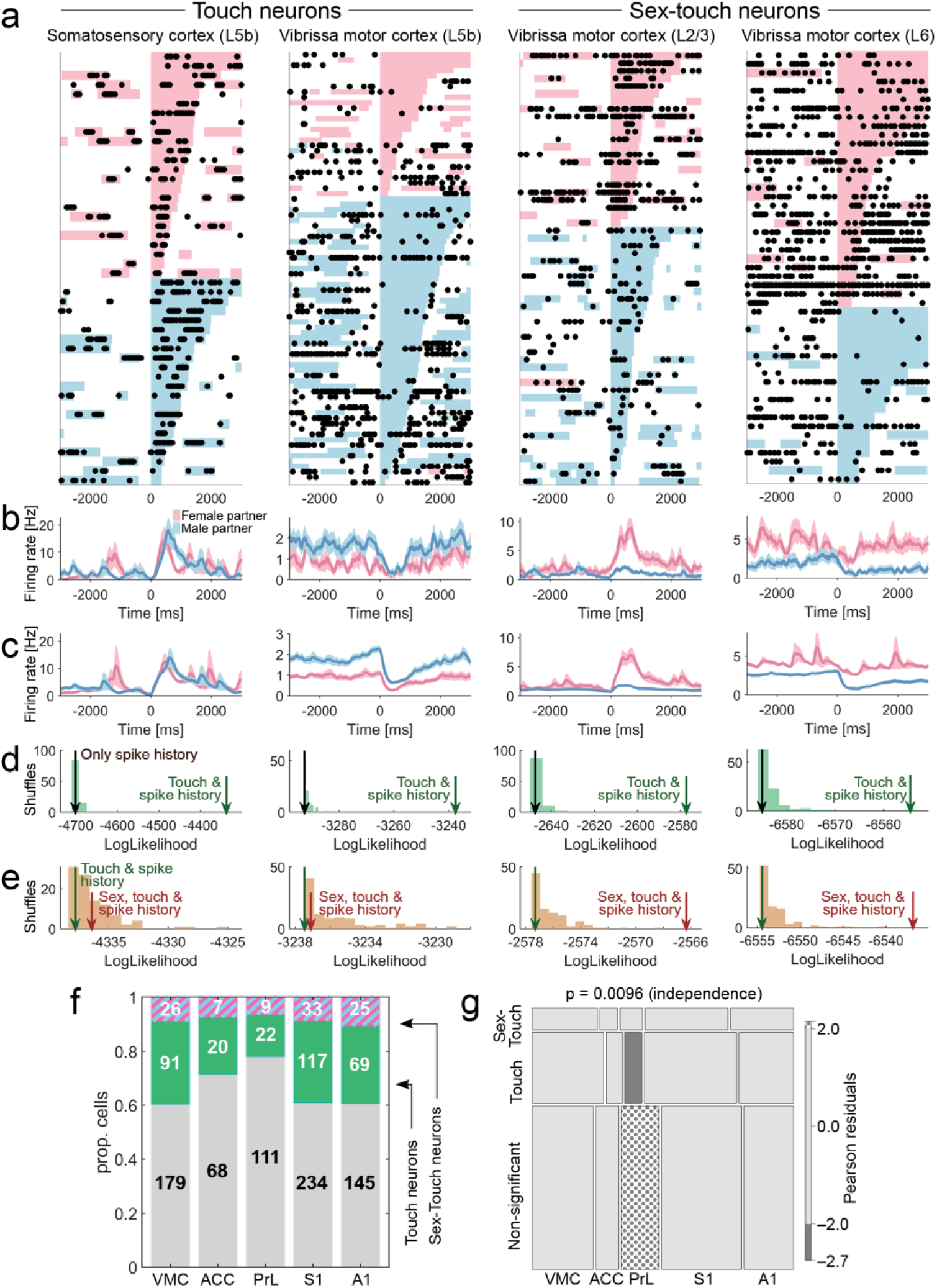
Single cortical neurons signal social touch and partner sex. **(a)** Raster plot of example touch (activated S1 L5b neuron and suppressed VMC L5b neuron) and sex-touch neurons (activated VMC L2/3 neuron and suppressed VMC L6 neuron). Raster plots show spike times (black dots) aligned to the first whisker-to-whisker touch in each social touch episode. Social touch episodes are sorted by partner sex (female: pink, male: blue) and by duration (indicated by length of colored bar). Many touch episodes happen close together in time and there is a large variability in the touch duration. (b) Peri-stimulus time histograms of the example neurons shown in (a), separated by partner sex. Line indicates mean firing rate, smoothed with an Alpha kernel (*τ* = 75 ms), shaded area indicates s.e.m, pink/blue color indicates female/male partner animals. (c) Peri-stimulus time histograms of the example neurons shown in (a), calculated from the fitted regression model, shown for comparison (plot conventions as in (b)).(d) Estimating touch-modulation: Log-likelihood values of models fitted to the neurons shown in (a). The log-likelihood of models depending on touch is indicated by the green arrow, the log-likelihood of models without touch is indicated by the grey arrow, and the log-likelihood distribution of shuffled touch-models is indicated by green bars. All neurons are significant at < 0.05 (the green arrow is outside shuffled distribution). (e) Estimating sex-touch-modulation: Log-likelihood values of models fitted to the neurons in (a). The log-likelihood of models depending on both partner sex and touch is indicated by brown arrow, the log-likelihood of models without sex is indicated by green arrow and the log-likelihood distribution of shuffled sex-touch-models is indicated by brown bars. The two touch neurons are not significantly modulated by sex (the brown arrows are inside shuffled distribution), both sex-touch neurons are significant at p < 0.05. (f) Number of neurons that modulated by touch (‘touch neurons’, green color) and neurons that are modulated by touch, but respond differently to male and female conspecifics (‘sex-touch neurons’, pink/blue striped color) across cortical areas. (g) Mosaic plot of the distribution of touch neurons, sex-touch neurons and non-significant neurons across cortical areas (p-value indicates χ^2^-test of independence). Colors indicate significantly increased (dotted) and decreased (grey) proportions (standardized Pearson residuals at p < 0.05).

As we expected based on the temporal patterns of the behavior (Figure 1e), a lot of the inter-trial variability in the baseline period was due to variations in behavior. For example, many of the spikes in the baseline period the example activated touch neuron (a layer 5b neuron in S1) and many of the pauses in firing in the example suppressed touch neuron (a layer 5b neuron in VMC) coincided with social touch episodes (Figure 3a). Similarly, much of the inter-trial variability in the post-stimulus the length of the social touch (Figure 3a).

Across all investigated areas, we found that a large proportion of neurons were modulated during social facial touch episodes. On average, across all areas, we found 25.3% touch neurons, and 8.3% sex-touch neurons (Figure 3f). We found that the proportions of touch and sex-touch neurons depended on the brain area (p = 0.0096, *χ*^2^-test of independence, Figure 2g) and a mosaic plot^37,38^ revealed that this was driven by the fact that PrL had less touch-neurons and more non-significant neurons than other brain areas (all: |standardized Pearson residual| > 1.96, all: p < 0.05, Figure 2g). This suggests that although there are differences in the proportions, information about touch and sex of the interaction partner is available across all investigated brain areas.

In a typical experiment, a subject animal would interact with at least two male and two female conspecifics (see Methods). In our analysis so far, we have naively grouped female and male interaction partners together and regressed to find neurons, that respond differently depending on the sex of the interaction partner (‘sex-touch’ neurons). We wondered if this observation was really due to the sex of the partner animal, or if partner-sex dependence could be an artifact of underlying individual-partner-specific responses. We used an information theoretical approach^39^ and found that even for the neurons most likely to have individual-specific responses, firing rates were significantly more informative about the actual sex than shuffled sex assignments, suggesting that biological sex – not individual partner identity – is the major determinant of touch responses (p = 0.007, paired model, Supplementary Figure 2).

### Structured population dynamics and depend on partner sex and subject sex

During social facial touch, different cortical regions have different prototypical response patterns^29,34^. Using our regression approach, we found that – as a population – SI neurons generally increased their firing rate during social touch, that VMC, ACC and A1 neurons generally decreased their firing rate and that PrL neurons displayed no modulation at the population level (all assessed at p < 0.05, Wilcoxon signed-rank tests, Supplementary Figure 3a). When we only analyzed neurons where touch modulation was significant at the single cell level, we found that these generally responded in the same direction as the population (Supplementary Figure 3b).

To investigate if the cerebral cortices of male and female rats process social touch differently, we first simply plotted all touch and sex-touch neurons recorded in the respective areas and separated neurons recorded in males and females (‘male neurons’ and ‘female neurons’). In SI, firing rates of both male and female neurons were increased by social touch and there was no difference between the sexes (p_male_, p_female_ < 0.001, male v. female p = 0.38, Figure 4a). In VMC, ACC and Al we found that both male and female neurons were generally decreased by touch, and we found no differences between the sexes there either (all: p_male_, p_female_ < 0.05, except p_female_ = 0.065 for ACC, p_male_ = 0.067 for Al, all male v. female p > 0.05, Figure 4a). In PrL, neither male nor female neurons were significantly modulated as a population, and there was no difference between the sexes (p_male_, p_female_ > 0.05, male v. female p > 0.05, Figure 4a).

**Figure 4.**
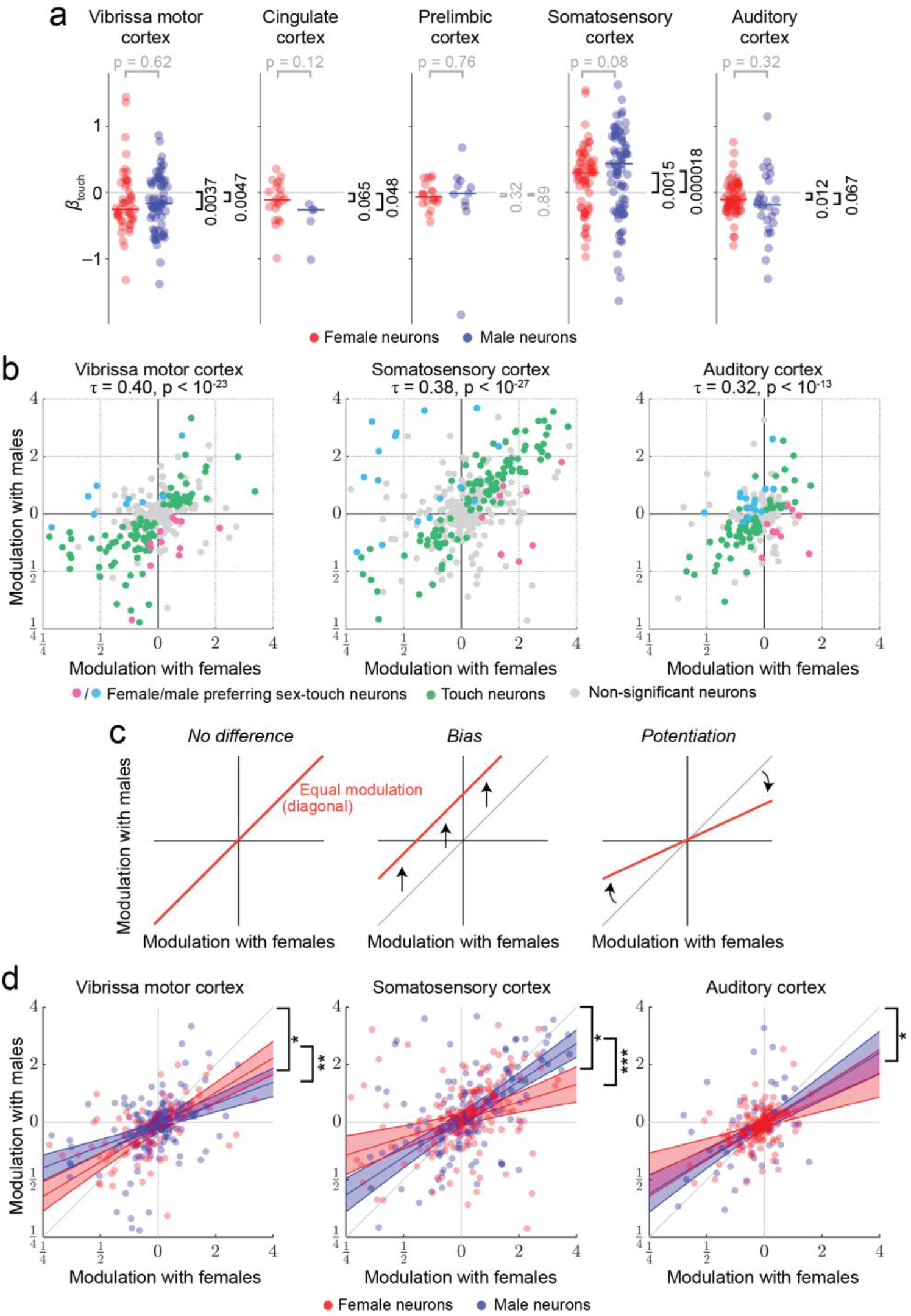
Population dynamics are highly structured and depend on both subject sex and partner sex. (a) Fitted *β*_*touch*_ of all touch and sex-touch neurons across cortical areas (axis clipped at +/−1.6, all data used for calculations, all data plotted in S2A-B). Colored dots indicate neurons recorded in female (red) and male (blue) animals. In S1, firing rates of both male and female neurons were increased by social touch (median male *β*_*touch*_ = 0.44, *p*_*male*_ = 0.000018, median female *β*_*touch*_ = 0.30, *p*_*female*_ = 0.0015, male v. female *p* = 0.08, N = 150). In VMC, ACC and A1, firing rates of both male and female neurons were generally decreased by touch (VMC: median male *β*_*touch*_ = −0.16, *p*_*male*_ = 0.0043, median female *β*_*touch*_ = −0.25, *p*_*female*_ = 0.0037, male v. female *p* = 0.62, N = 117, ACC: median male *β*_*touch*_ = −0.26, *p*_*male*_ = 0.048, median female *β*_*touch*_ = −0.11, *p*_*female*_ = 0.065, male v. female *p* = 0.12, N = 27, A1: median male *β*_*touch*_ = −0.18, *p*_*male*_ = 0.067, median female *β*_*touch*_ = −0.10, p_female_ = 0.012, male v. female p = 0.32, N = 94). In PrL, neither male nor female neurons were significantly modulated as a population, and there was no difference between the sexes (median male *β*_*touch*_ = −0.015, *p*_*male*_ = 0.89, median female *β*_*touch*_ = −0.067, p_female_ = 0.32, male v. female *p* = 0.76, N = 31). All: t-tests (*p*_*female*_, *p*_*female*_) and unpaired *t-test* with unequal variance (male v. female) if normal by a Lilliefors test, else Wilcoxon signed rank tests and Mann-Whitney U-test, respectively. * indicates p < 0.05, ** indicates p < 0.01, *** indicates p < 0.001. (b) Modulation of activity (in fold change) during social touch with male and female conspecifics is highly correlated (VMC/S1/A1: Kendall’s *τ* = 0.40/0.38/0.32, all *p* < 10^−23^/10^−27^/10^−13^). Touch neurons are indicated by green dots, female/male preferring sex-touch neurons are indicated by pink/blue dots and non-significant neurons are indicated by grey dots, Kendall’s *τ* and p-value above. (c) Some possible types of population structure. If there is no overall difference between responses to male and female conspecifics, neurons will fall on the diagonal (left). A biased response to one partner sex corresponds to a shift away from the diagonal. For example, the red line corresponds to a situation where neuronal responses to female conspecifics are always larger in magnitude than responses to male conspecifics (left). (d) Population response pattern depends on subject sex and partner sex. Dots indicate neurons recorded in female (red) and male (blue) subject animals, lines indicate maximum-likelihood fit of regressing modulation with males as a function of modulation with females (red = female, blue = male, shaded area indicates 95% C.I.) * indicates slope different from unity (outside 95% C.I. for both males and females), ** indicates p < 0.01, *** indicates p < 0.001 (full model specification in Suppl. Note 2).

Next, we investigated if population responses depend on the sex of the interaction partner. In the posterior parietal cortex (PPC), there is evidence that modulation by sensory stimuli and movement features is distributed randomly across neurons with no or little population structure^40^. To understand if the same is true during social touch, we plotted the modulation of neurons during interactions with male and female conspecifics. Similar to the PPC, we found that responses were veiy diverse and that neurons populated all quadrants (Figure 4b, Supplementary Figure 4). Sex-touch neurons were both in the first and third quadrants (corresponding to differences in magnitude of modulation) and second and fourth quadrants (corresponding to different directions of modulation, e.g. increasing during interactions with males and decreasing during interactions with females). As expected from the definition, touch neurons were generally near the diagonal and non-significant neurons were near the origin (Figure 4b).

First, we focused on the three brain areas where we had a substantial dataset (VMC, SI and Al). In these areas, we found that responses to male and female interaction partners were highly correlated (VMC/S1/A1: Kendall’s *τ* = 0.40/0.37/0.32, all p < 0.001, Figure 4b). The high correlation values suggest that the population response to male and female stimuli is nonrandom. But what is the structure of the population response, and does it depend on the sex of the subject animal? High correlation values could reflect that there is actually no difference between responses to males and females. This would correspond to having all neurons along the diagonal, and could mean that the sex-touch neurons identified above might simply be neurons at the edge of this unity-distribution (Figure 4c, left). If responses to males and females are different, the responses could be biased towards one sex, responses when touching one sex could be potentiated, or it could be a combination of both. A biased response would for example correspond to having all neurons spike more when interacting with males than females (Figure 4c, middle). A potentiated response would for example mean that responses when touching females are always stronger in magnitude than responses when touching males (Figure 4c, right). Finally, these response patterns might depend on the sex of the subject animal.

To identify the population pattern, we performed mixed-effect generalized linear modeling to investigate the how responses to male and female partners depend on the partner-sex-by-subject-sex contingency (see Methods). We did not find any evidence of a bias in responses (all: *p*_*intercept*_ > 0.05) and no subject-sex-dependent bias (all: *p*_*subject_sex*_ > 0.05, full model specification in Supplementary Note 2). However, in all three areas, we found evidence of potentiation. There was a highly significant dependence of responses to males on the response to females (all: *p*_*subject_sex*_ < 0.001) and the estimate of the slope was significantly smaller than unity, suggesting that responses to females are stronger in magnitude than responses to males (VMC/S1/A1: βsubject_sex [C.I.] = [0.27-0.49]/[0.54-0.78]/[0.45-0.78], Figure 4d). In SI and VMC, there was also a significant interaction between the subject sex and this potentiation (both: *P*_*female_modulation*subject_sex*_ < 0.01, Suppl. Note 2). Interestingly, the interaction had opposite signs (VMC/S1: *β*_*female_modulation*subject_sex*_ [C.I.] = [0.055-0.39]/[−0.54,−0.16], Figure 4d).

We also found significant correlation values between responses to male and female conspecifics in the areas where we only had a limited sample (ACC/PrL: Kendall’s = 0.22/0.17, both p < 0.01, Supplementary Figure 5a). In these smaller datasets, there was no statistically significant subject-sex-dependent or stimulus-sex-dependent potentiation, but – although not significant – the maximal-likelihood fit also estimated (*β*subject_sex to be less than unity for both ACC and PrL (Supplementary Figure 5b, Supplementary Note 2). We note that the fact that the pattern is similar in the areas with little data, might suggest that this partner-sex-dependent potentiation is very wide-spread across cortex. We also note that in ACC we did see a significant bias, indicating that ACC neurons fire less with male than female conspecifics, in both male and female subjects (pintercept < 0.05, Supplementary Note 2).

In summary, for both male and female subjects, cortical responses were larger in magnitude with female than male interaction partners (statistically significant in VMC, SI and Al, same direction, but not significant in ACC or PrL). In SI, the potentiation of responses to females was stronger in female subjects than male subjects; in VMC, the potentiation was stronger in male subjects; and in Al, the potentiation did not depend on the subject sex.

### Plausible cellular mechanisms underlying population responses

We found that both partner sex and subject sex was encoded in a potentiation of social touch responses. This potentiation pattern suggests that a potential mechanism could be a change in inhibitory drive, similar to how inhibitory neuron subtypes control context-dependent modulation of sensory responses in visual^41,42^ and auditory cortex^43^ (for review see ref. ^44^). A likely candidate mechanism for the regulation of cortical interneuron activity during social touch is by neuro-modulatory hormones, such as estrogen and oxytocin, both key regulators of sociosexual behavior.

While classically thought to mainly act on slow time-scales (~hours), estrogen can be rapidly synthesized in the brain and might affect sensory processing at much faster time scales (< min)^45^. We have previously reported that baseline firing rates in putative excitatory neurons in rat somatosensory cortex is lower during estrus^29^, that deep layer parvalbumin-positive interneurons in rat somatosensory cortex express estrogen receptor β and that fast-spiking interneuron activity increases in estrus and with estradiol injection^31^. Despite the change in baseline firing rate, however, we previously reported social-touch responses during staged social interactions in head-fixed animals were unchanged across the estrus cycle, both for in regular-spiking and fast-spiking neurons^31^. In our dataset, we have access to the estrus state of female subject animals in a subset of the data (only from somatosensoiy cortex) and in these data, we also found that responses to social touch were unchanged across the estrus cycle (Supplementary Figure 6).

Oxytocin action on neural circuits plays a major role in cognitive and emotional aspects of sociosexual behavior, such as social bonding, anxiety and trust^46,47^ and intranasal oxytocin affects the subjective pleasantness of identical touch stimuli^13,19^. Oxytocin receptors are expressed in cortex mainly by interneurons^48,49^ and oxytocin acts on interneurons in auditoiy cortex to enable pup-retrieval by mother mice^50^ and modulates interneuron activity in prefrontal cortex^49^ and olfactory cortex^51^ to enable social recognition. Gentle touch and stroking activates oxytocinergic neurons in the paraventricular hypothalamus^52^ which project directly to sensoiy cortices^48^. This provides a potential circuit, by which touch-related oxytocin release can impact cortical interneuron activity to modify cortical responses and network activity patterns during social facial touch. Moreover, since the pattern of cortical oxytocin receptor expression is sex-depcndcnt^48,53,54^, this provides a potential basis for our observed sex differences in network activity patterns^55^.

In order to determine how transient oxytocin release might affect cortical population responses during social touch, we simulated the effect of oxytocin release using a large-scale spiking neural network model of somatosensoiy cortex* (Supplementary Figure 5). This model allows us to simulate the activity of ~1 mm2 of somatosensory cortex (~80.000 modeled neurons, with ~300 million synapses) with realistic synaptic connectivity and projection patterns based on functional and anatomical studies^56^ (see Methods). This model has independently validated multiple times and reproduces cell-type-specific and layer-specific distributions of firing rates, correlations and spike delays^56–59^.

We still know little about how exactly oxytocin modulates neural processing across cortical regions. Interestingly, however, the little that we do know paints a cohesive picture: Multiple recent studies investigating oxytocin action in both neocortex and isocortex (hippocampus) have observed two main effects, one on interneurons and one on excitatory neurons: (i) Increased firing rate of inhibitoiy interneurons, with no change in IPSC amplitude, likely due to membrane depolarization^46,48–50,60^ (ii) Increased excitability of excitatoiy neurons, visible as an increase in input resistance^61,62^.

In our simulations, we thus investigated how a transient depolarization of interneurons and/or a transient increase in the input resistance of excitatoiy neurons modulated the population response to ‘touch’ (i.e. thalamic input, two example ‘touch’ trials shown in Figure 5b). We made three observations. First, we found that increasing inhibitoiy drive by depolarizing interneurons lead to a potentiation of the population response, the same type of modulation as we observed during social touch (Figure 5c, first column: the slope is different from unity, and there is no shift in the response, compare with Figure 4b-d). Second, we found that increasing the input resistance of excitatoiy neurons simply lead to a bias in responses, not what we observed during social touch (Figure 5c, first row: there is a constant upwards shift, and the slope is not different from unity). Third, we found that when both modulations were simulated simultaneously, the effects “cancel out” and the population response is approximately normalized (Figure 5c, on the diagonal: the slope is unity with only a very slight bias, intermediate configurations visible in the off-diagonal panels).

**Figure 5.**
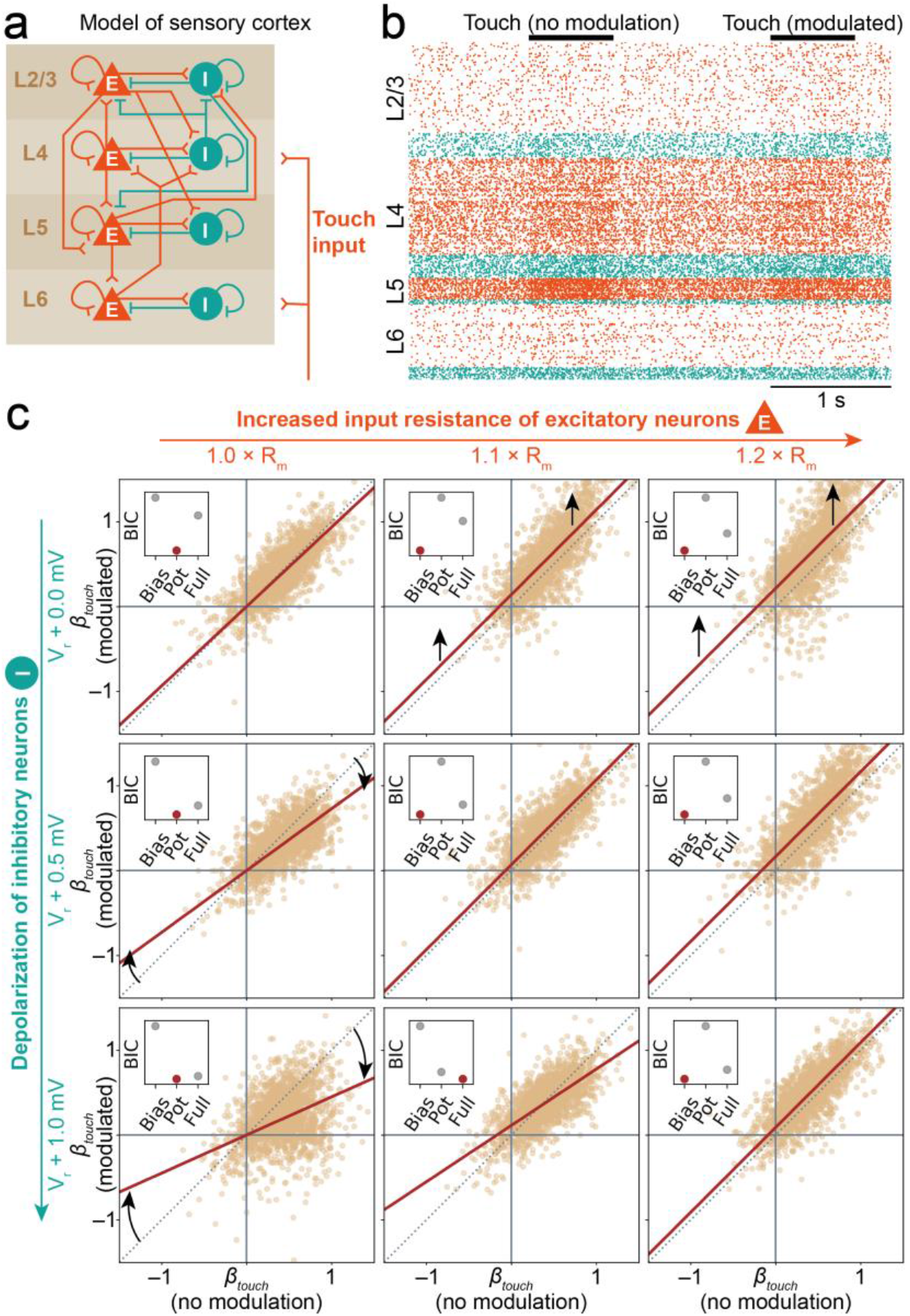
Modulation of cortical inhibition could underlie population dynamics during social touch. (a) We simulated the activity of ~1 mm^2^ of sensory cortex as 77,169 leaky integrate-and-fire neurons with biologically realistic cell-type specific connectivity (~0.3 billion synapses)^56,59^. The model consists of four cortical layers (2/3, 4, 5 and 6), each with a population of excitatory (“E”, orange pyramids) and inhibitory (“I”, teal circles) neurons. A population of simulated thalamic neurons provides excitatory synaptic input to layers 4 and 6 during simulated touch. This graphical representation of the model displays the major excitatory and inhibitory projections (only connection probabilities > 0.04 are drawn, full connectivity matrix shown in Methods). (b) Raster plot showing activity of a random subset (2%) of the modeled neurons, during two simulated touch trials (each row is a single neuron, orange/teal dots indicate the spike times of excitatory/inhibitory neurons, neurons are sorted by layer). In our simulations, we simulated ‘touch trials’ as 1000 ms of spontaneous activity with no thalamic input, followed by 700 ms of activity with thalamic input (‘Touch’, indicated by black bars), followed by 300 ms of activity with no thalamic input. ‘Non-modulated’ and ‘modulated’ trials were interleaved. (c) Simulated population dynamics, when we simulate neuromodulation by oxytocin release during social touch. When we do not modulate the network during touch (V_r_ + 0.0 mV, 1.0 × R_m_, top left corner) the responses during ‘modulated’ touch and ‘nonmodulated’ touch is the same (reset voltage of the interneuron populations, V_r_, and membrane resistance of the excitatory popluations, R_m_, indicated outside plots, each dot indicates the touch response of 2000 single random neurons, brown line shows the best model, small inserted plot indicates the Bayesian information criterion of three models fitted to the data: a ‘Bias model’ with only a bias and the slope fixed at unity; a ‘Potentiation model’ with no bias, but a free slope parameter; and a ‘Full model’ allowing for both a bias and a potentiation of responses, compare with Figure 4b). The simulations invite three conclusions: Increasing inhibitory drive by depolarizing interneurons lead to a potentiation of touch responses, just as we observed in our data (first column: slope different from unity, no shift. Compare with Figure 4b-d). Increasing the input resistance of excitatory neurons simply lead to a bias in responses (first row: upwards shift from unity line). When both effects are applied simultaneously, the effects “cancel out”, and the touch response is approximately normalized (diagonal: the best model again essentially falls on the unity line).

As a low-level validation of the modeling approach, we compared the responses of simulated excitatoiy and inhibitory neurons to the response of putatively inhibitoiy and excitatoiy neurons in our somatosensoiy cortex data (assessed by spike shape, see Methods). In real data from SI (Supplementary Figure 7b-d), as well as in data from Al and VMC (Supplementary Figure 7e-j), we found that responses of putatively inhibitoiy and excitatory neurons were similar, and in the same direction. This overlap in response magnitudes, and the similar direction of firing rate modulation of agrees with the prediction of the model (Supplementary Figure 7a).

From these simulations, we draw two conclusions. First, we conclude that changes in inhibitory drive – for example mediated by oxytocin release – is a plausible candidate mechanism, which could explain the population response potentiation during social facial touch. Second, as an additional incidental finding, we reason that the increased excitability of excitatory neurons that is observed after sustained bath application of oxytocin^61,62^ might be a homeostatic response which “re-normalizes” network processing in answer to oxytocin-mediated increases in inhibition during such in-vitro experiments.

## Discussion

A previous study found that firing rate modulation by sensory stimuli and movement features in the posterior parietal cortex was distributed randomly across neurons with no or little population structure^40^. In contrast to this observation, we found that social touch responses were highly structured, and that the network activity pattern signaled the sex of the interaction partner by partner-sex-dependent potentiation of responses. Moreover, in somatosensoiy and vibrissa motor cortex we found that the sex of the subject animal itself was encoded by a subject-sex-dependent magnitude of this potentiation. In line with the encoding of partner sex observed, there is previous evidence of population coding of touch stimuli in the somatosensory cortex^63,64^. Previous investigations in the rodent whisker system have found that putatively ‘simpler’ stimuli presented in highly controlled conditions, such as object location^65,66^ and texture roughness^67^, are encoded with higher fidelity by precise spike timing than by firing rates. However, in line with our observations, previous investigations also found that firing rates cany extra information in addition to the information conveyed by spike timing^68–70^. The relevant time scale for temporal coding by spike timing in the somatosensory cortex seems to be within tens of milliseconds from the stimulus, and the analysis requires extremely precise information about stimulus onset time^66,71,72^, which is not available in our naturalistic experimental paradigm. More generally, structured population dynamics in vibrissa motor cortex aligns well with other observations of population coding in motor cortex, where the population activity vector of both preparatory^73^ and movement-related activity^74 75^ correlates with movement features^76–78^.

In line with our observations of touch- and sex-touch neurons in rat cingulate cortex, previous human studies have found that social context strongly modulates responses in cingulate and orbitofrontal cortex^18–20^. We found that – when examined at single cell resolution – the majority of cingulate neurons decrease in activity during social touch (a positive social interaction). This is in stark contrast to a recent report that only ~1% of rat cingulate recording sites show decreases in multiunit activity during a painful stimulus, during playback of a fear-conditioned tones or when observing conspecifics receive painful electric shocks^79^. In light of the tight anatomical integration of prelimbic cortex with brain structures responsible for social behavior and emotions^80,81^ and the putative homology with human and primate prefrontal cortex, a key structure in social cognition^82,83^, we found it curious that prelimbic cortex was so weakly modulated by social touch. We did not see any overall changes in firing rate, only weakly modulated touch and sex-touch neurons at both edges of a zero-centered distribution. While previous rodent studies did not investigate social touch as such, activity of subpopulation of prelimbic neurons are modulated by the presence of conspecifics^84^, but – in line with our observations – it has been shown that during social approach (to a caged conspecific) prelimbic responses are generally weak and diverse^85,86^ and that social behaviors are encoded by sparse groups of neurons either increasing or decreasing in activity during behavior (‘on’ and ‘off’ ensembles^87^).

In line with what we described for other whisking behaviors^34^, we found that despite active whisker movement, neurons in vibrissa motor cortex mainly decrease their activity during social touch. We also found a large proportion of sex-touch neurons in vibrissa motor cortex. We already know that neurons in rodent motor cortex can have low-latency responses to touch^88–93^, that motor cortex is an important node in a distributed network for touch processing^94–98^ and that motor cortical activity modulates somatosensory processing in both somatosensory cortex^99–101^ and thalamus^102^. Vibrissa motor cortex has been implicated in diverse aspects of sensorimotor cognition (for reviews, see ^103,104^) and our study adds a potential role in sex-dependent processing of social cues to this complex picture.

In somatosensory and motor cortex, we found that social touch leads to different network activity patterns in male and female subjects. Identifying such potential differences in how male and females cortices process social stimuli could help shed light on the biological basis of sex differences in the etiology, prevalence and symptoms of autism, depression and anxiety disorders, all characterized by social dysfunction^105–107^. Generally, while there clearly are some systematic sex differences in both anatomy^108–113^ and functional connectivity^114^, male and female cortices are overall remarkably similar. For example, even though male and female genital anatomy is extremely different, the layout of the somatosensory body map is essentially identical in both sexes^115,116^ and has similar projection targets^117^. In recent years, several studies have identified sex differences in sub-cortical circuits involved in regulating social behavior. For example, circuits involved in the control of aggression in the ventromedial hypothalamus are wired differently in males and females^118^, medial amygdala has striking sex differences in olfactory responses^119^, and galanin-positive neurons in the medial preoptic area^120^ and their subcortical input nuclei^121^ are activated during parental behavior in a sex- and reproductive-state-specific manner. In contrast to these sub-cortical examples, sex-differences in cortical processing are rare and subtle^83,122,123^.

Our data invites two, not mutually exclusive, interpretations. On one hand, the differences in processing may be a signature of a universal sex difference in how male and female cortices process the same sensory stimuli. Thus, these sex differences might generalize across other cortical regions and sensorimotor modalities. On the other hand, male and female cortices may use identical computational strategies, and our observed differences simply reflect the fact that for male and female subjects, the same partner sex presents a different social situation with different cognitive and behavioral challenges. For example, a male rat interacting with another male rat might be mainly assessing dominance and aggression, while a female rat meeting a male rat might be assessing aggression as well as reproduction^27,124^.

We still know very little about sex differences in human cortical processing of social touch stimuli. As noted in the introduction, our investigation of partner-sex dependent differences in touch responses was based on pioneering studies with human subjects^18–20^. These studies have not reported sex differences in the processing of social touch stimuli, but then again, previous studies have not compared experimental subjects of both sexes^13,16–21,125–127^. It would be most interesting to know if our observed network activity differences in the rat are paralleled in primates and humans. Our findings highlight the importance of complementing human work with the single cell resolution offered by animal studies.

During social facial interactions, the actual haptic input from male and female partners is very similar^27^. However, our naturalistic paradigm is inherently multisensory, and even though the whisking and touch input is similar, male and female partners convey very dissimilar olfactory cues. The vomeronasal organ is important for determining the sex of the conspecific in both male-male^128^ and male-female interactions^129^, and might be an important ‘bottom-up’ pathway for the modulation of cortical responses during social touch observed here. Still, other sensory modalities might also contribute. For example, there might be subtle sex-differences in whisking patterns^27^ or vocalization patterns. We know that rats vocalize around and during social facial interactions^33^ and that rat pups^130^ and adult mice^131^, for example, display subtle sex differences in vocalizations. Our definition of social ‘touch’ neurons is purely descriptive and we do not know what low-level features of the social touch episode (touch, ultrasonic calls, olfactory cues, temperature) or internal ‘top-down’ processes modulate the firing rate of these neurons.

We found that changes in social context (partner sex and subject sex) were associated with a structured potentiation of social touch responses. The neuromodulatory hormone oxytocin is a prime candidate for cortical neuromodulation during social interactions^13,46,47,132,133^ and our simulations suggest that changes in inhibitory activity by social-touch-dependent release of oxytocin could plausibly underlie the observed context-dependent modulation of touch responses. Across all areas, we observed that responses to female partners tended to be bigger in magnitude than responses to male partners. Thus, if the touch response potentiation is indeed mediated by oxytocin using the simple mechanisms outlined in our simulations, this makes the prediction that oxytocin release should be larger when interacting with a male than a female partner. Does this prediction match what has been reported in the literature? Surprisingly, there is a real lack of comparative studies between males and females^54,132^, and we could only find two reports which speak to this question. When heterosexual women touch their male partners, they release more systemic oxytocin than heterosexual men do when touching their female partners^134^, and male rats release more systemic oxytocin when socially interacting with male partners than female rats do when socially interacting with female partners^135^. These two observations are obviously consistent with our prediction, but they constitute extremely weak and indirect evidence. We still know very little about what drives cortical oxytocin release on a moment-to-moment basis during social interactions, and whether cortical oxytocin release depends on the sex of the interaction partner^46,47,132,133^.

In any case, it is important to stress that neuromodulation by oxytocin is just one among many possible mechanisms. Meeting a whisking, vocalizing and phero-mone-scented conspecific en face is presumably a highly salient event and the response modulation across cortical networks could reflect a generalized change in cortical ‘state’ (alertness, elicited by the high stimulus salience), perhaps to increase the signal-to-noise of sensory processing^136–138^. The social-context-dependent changes in touch responses observed during social facial touch might be partly or fully mediated by other neuromodulators, such as vasopressin, another peptide hormone involved in the control of social behavior^47,133,139^; dopamine, which might strongly modulate cortical activity during social motivation and reward^140–144^; or acetylcholine or noradrenaline, both known to modulate cortical state and cortical sensory processing in a context-dependent manner^44,138,145,146^.

## Methods

### Animal welfare

All experimental procedures were performed according to German animal welfare law under the supervision of local ethics committees. Rats (Wistar) were purchased from Janvier Labs (Le Genest-Saint-Isle, France). Rats presented as partner animals were housed socially in same-sex cages, and postsurgery implanted animals were housed in single animal cages. All animals were kept on a 12h:12h reversed light/dark cycle and all experiments were performed in the animals’ dark phase. Rats had ad libitum access to food and water.

### Social gap paradigm

Behavioral experiments were done using the social gap paradigm^27^ (Figure 1a). The experimental paradigm consists of two elevated platforms, 30 cm long and 25 cm wide surrounded by walls on 3 sides, positioned approximately 20 cm apart. The distance between platforms was varied slightly depending on the size of the rats. The platforms and platform walls were covered with soft black foam mats to provide a dark and non-reflective background and to reduce mechanical artifacts in tetrode recordings. All experiments were performed in darkness or in dim light, and behavior was recorded from above under infrared light. The implanted rat was placed on one platform, and on the other platform we either presented various objects or other rats. The implanted rats were not trained, just habituated to the setup and room, and spontaneously engaged in social facial interactions. The rat behavior was recorded at low video speed from above with a 25 Fiz or 30 Fiz digital camera, synchronized to the electrophysiological data acquisition using TTL pulses. Typically, recording sessions were performed in four to eight 10-15 min blocks, where we would present either one or two conspecifics (of both sexes) in each block, randomly, see^29,33,34^. Video frames were labeled blind to the spike data.

### Tetrode recordings and histology

In tetrode recording experiments, we used ~p60 Wistar rats, which were handled for 2-3 days before being implanted with a tetrode microdrive. Surgery was done as previously described^29^. The implanted microdrive had eight separately movable tetrodes driven by screw microdrives (Harlan 8-drive; Neuralynx, Bozeman, MT, USA). The tetrodes were twisted from 12.5 μm diameter nichrome wire coated with polymide (California Fine Wire Company), cut and examined for quality using light microscopy and gold-plated to a resistance of ca. 300 kOhm in the gold-plating solution using an automatic plating protocol (“nanoZ”, Neuralynx). For tetrode recordings targeting VMC, ACC and PrL, a craniotomy of 1×2 mm was made 0.75-2.75 mm anterior and 1-2 mm lateral to bregma^147,148^. ACC includes data from the area ‘Cgl’ and dorsal ‘Cg2’ in the atlas^147^. A1 includes data recorded in the rat atlas regions Aul’, AuD’ and AuV’^147^. Steel screws for stability and two gold-plated screws for grounding the headstage were drilled and inserted into the skull, and the gold-plated screws were soldered and connected to the headstage PCB using silver wire. After fixation of all screws, the dura was removed, the implant fixated above the craniotomy, the craniotomy sealed with 0.5% agarose and the tetrode drive fixed in place with dental cement (Heraeus). The tetrodes were arranged in a 2-by-4 grid with a spacing of ~500 μm. Neural signals were recorded through a unity-gain headstage preamplifier and transmitted via a soft tether cable to a digital amplifier and A/D converter (Digital Lynx SX; Neuralynx) at 32 kHz. We filtered the signal between 600 Hz and 6 kHz and detected spikes by crossing of a threshold (typically ~50 μV) and saved each spike (23 samples, 250 μs before voltage peak to 750 μs after voltage peak). At the end of the experiment, animals were again anaesthetized with a mix of ketamine and xylazine, and the single tetrode tracks were labelled using small electrolytic lesions made by injecting current through the tetrode wire (10 μA for 10 s, tip-negative DC). After lesioning, animals were perfused with phosphate buffer followed by a 4% paraformaldehyde solution (PFA). Brains were stored overnight in 4% PFA before preparing 150 μm coronal sections. Sections were stained for cytochrome oxidase to reveal the areal and laminar location of tetrode recording sites, which could be calculated from the location of tetrode tracks and lesions. We only analyzed data from recording sites where the lesion pattern could unanimously identify the tetrode and the recording sites.

All spike analysis was done in Matlab (MathWorks, Natick, MA, USA). Spikes were preclustered off-line on the basis of their amplitude and principal components by means of a semiautomatic clustering algorithm (‘KlustaKwik’, https://github.com/klusta-team/klustakwik, ^149^). After preclustering, the cluster quality was assessed and the clustering refined manually using MClust (http://redishlab.neuroscience.umn.edu/MClust/MClust.html, A. D. Redish, University of Minnesota). The spike features used for clustering were energy and the first principle component of the waveform. To be included in the analysis as a single unit, clusters had to fulfill the following criteria: first, the L-ratio, a measure of distance between clusters, was below 0.5. Second, the histogram of interspike intervals (ISIs) had to have a shape indicating the presence of single units, e.g. a refractory time of 1-2 ms, or the appearance of a bursty cell (many short ISIs). Multi-unit clusters were not included in the analysis.

### Previous use of data

Part of the data presented in this study has already been presented in other studies. The barrel cortex data, recorded throughout all cortical layers, has previously been published in ref. ^29^. The auditory cortex data, recorded throughout all cortical layers, has previously been published in ref. ^33^. The recordings from deep layers of vibrissa motor cortex has previously been presented in ref. ^34^. The previous study on barrel cortex’’ investigated how touch-evoked activity depended on male and female subject and partner animals in a PSTH-based way (like our Figure 1C-D), the other two previous studies did not investigate any sex differences^33,34^. Neuons were recorded throughout the cortical column, with a majority of neurons in the deep layers. The laminar distribution was (Sl/VMC/PrL/ACC/Al): Layer 6: 32/16/39/31/46, layer 5b: 88/153/70/17/100, layer 5a: 42/37/33/47/63, layer 4: 56/0/0/0/19, layer 2/3: 37/90/0/0/4, layer uncertain: 129/0/0/0/7. All data from the superficial layers of vibrissa motor cortex, data from all layers of cingulate cortex and data from all layers of prelimbic cortex have not previously been published.

### Statistical modeling – PSTHs

To identify significantly increasing and decreasing neurons, as shown in Figure 1C-D, we calculated the mean firing rate in the ‘baseline’ period (2500 to 0 ms) and the post-stimulus period (0 – 500 ms) and determined significant changes using Wilcoxon signed-rank tests. For display, the PSTHs were smoothed with PSP-shaped alpha function (i.e. 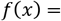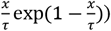) with *τ* = 75 ms.

### Statistical modeling – Touch and sex-touch neurons

We used a generalized linear regression approach^35,36^ to identify touch and sex-touch responses. We discretize the spike train in 1-ms bins, reduced the data amount by removing baseline periods, which were more than five seconds from any social interaction and model the firing rate as a Poisson process. If we assume that the discharge of spikes within each time bin is generated by a homogenous Poisson point process, then the probability of observing y spikes in a single time bin is

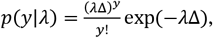

where Δ = 1 ms is the width of the time bin and λ > 0 s^−1^ is the expected discharge rate of the cell. If we assume that each time bin is independent, the prohability of the entire spike train, 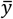 is

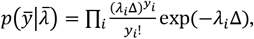

where *y*_*i*_, λ_*i*_ is the observed number of spikes and the expected discharge rate in the i’th time bin, respectively. If we model the expected discharge rate, 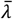, so that it depends on the parameters, 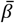, we have the log-lilcelihood function

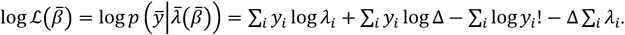

We model 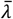 so that it depends linearly on spike histoiy, experimental recording, touch and partner sex and – since the expected firing rate cannot be negative – we model

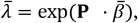

where P is a predictor matrix and 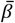 is a vector of regression coefficients ^35^. The predictor matrix has the following columns: a constant baseline rate; five 1-ms spike history bins; six 25-ms history bins; (*N*_*rec*_ − 1) one-hot columns to model a change in baseline between the recordings; a one-hot column indicating all social touch episodes; a one column indicating the sex of the stimulus animal (0 = female, 1 = male). Due to the refractory period of the cell, it is not correct to assume that all time bins are statistically independent, so following we include 11 spike history parameters, h_1_…h_11_, to model the inter-spike-interval distribution of the cell. The spike history is binned to 11 successive bins, five 1-ms bins (vectors with no. of spikes in the previous 0-1 ms, 1-2 ms, 2-3 ms, 3-4 ms, 4-5 ms) and six 25-ms bins (vectors with no. of spikes in the previous 5-30 ms, 30-55 ms, 55-80 ms, 80-105 ms, 105-130 ms, 130-155 ms). We also include constant bias terms to allow for variations in baseline firing rate between each recording. Thus,

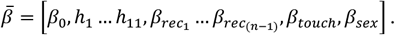

and in Wilkinson notation, the linear model would be written,

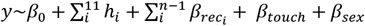

An example predictor matrix is shown in Figure S1A. We used the function package ‘neuroGLM’ (https://github.com/pillowlab/neuroGLM, ref. ^150^) to calculate and numerically fit the models using Matlab (MathWorks, Natick, MA, USA). To assess the statistical significance of touch and sex-touch responses, we used a non-parametric, shuffling-based model selection approach. To assign statistical significance to *β*_*sex*_, we compared the log-likelihood of the model including touch and sex as predictors with the distribution of log-likelihood values from the same models fitted to predictor matrices, where we randomly shuffled the label (male/female) of the partner animal (N = 100, Figure 3D). We do not assume that random effects have a Gaussian distribution. Rather, to assign statistical significance to *β*_*touch*_, we compared the log-likelihood of the model including touch (and not sex) as predictors with the distribution of log-likelihood values from the same models fitted to predictor matrices, where we circularly permutated the column indicating where the social touch episodes had happened (N = 100, Figure 3E). We defined ‘sex-touch neurons’ as neurons with *p*_*sex*_ < 0.05, ‘touch neurons’ as neurons with *p*_*sex*_ > 0.05 and *p*_*touch*_ < 0.05, and ‘non-significant’ neurons as neurons with both *p*_*sex*_ > 0.05 and *p*_*sex*_ > 0.05. We compared the proportions of non-significant, touch and sex-touch neurons across cortical areas by calculating standardized Pearson residuals and visualized the contingency table as a mosaic plot^37,38^. Standardized residuals and mosaic plots were generated using the ‘vcd’ package^151^ for R^152^.

### Information theory

To avoid assuming anything about the direction of firing rate changes, we calculated the information per spike,

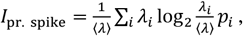

where ⟨λ⟩ is the average spike rate, λ_*i*_ is the spike rate during the i’th stimulus, and and p, is the probability of the i’th stimulus^39^. To identify neurons, which could potentially have individual-specific or sex-specific response patterns we treated as ‘stimuli’ the social touch episodes with all individual partner animals. Significance was assessed by a shuffling procedure, where we circularly shifted the timing of the social touch episodes over the recordings (N = 200, p<0.05).To investigate if these putatively individual-specific neurons carry more information about the real sex of the partner animal than randomly assigned sex labels, we also used a shuffling procedure. For all neurons, we calculated the mutual information per spike with three situations, baseline, social touch with male partners and social touch with female partners, and calculated the same value, where we shuffled the sex of the partner animals (Figure S2A). Since the number of possible shuffles depend on the number of partner animals, the dataset had strong dependency per neuron^153^, so we fitted a mixed effects model,

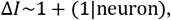

where Δ*I* is the real minus the shuffled value of the information per spike between the three ‘stimuli’.

### Statistical modeling – Magnitude of responses

To estimate the average depth of modulation of increasing and decreasing neurons across both partner sexes (Figure 3), we calculated the fold modulation bv touch as the ratio

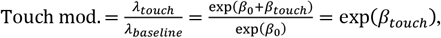

where *λ*_*touch*_, *λ*_*baseline*_ are the firing rates and *β*_*touch*_ is the fitted regression coefficient of the GLM models including touch (and not sex) as a predictor. To plot and compare the magnitude of increases and decreases, we calculated the numerical value of the base-2 logarithm and plotted the data as fold increases/decreases (e.g. log_2_ (ratio) = 1 corresponds to a two-fold increase (double the firing rate), log_2_ (ratio) = −1 corresponds to a two-fold decrease (half the firing rate), etc.). To estimate the modulation when touching male and female conspecifics (Figure 4), we calculated the fold modulation from the fitted GLM models as

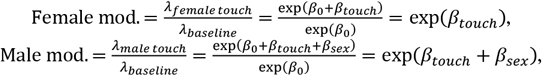

where *λ*_*female touch*_, *λ*_*male touch*_,*λ*_*baseline*_ are the firing rates and *β*_*touch*_ *β*_*sex*_ are the fitted regression coefficients of the full GLM models including touch and sex (i.e. this is a different fitted value of *β*_*touch*_ than above, Touch mod. ≠ Female mod.). To determine the population response pattern of the modulation, we fitted the mixed-effects regression

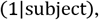

where male_mod and female_mod are the modulation ratios, subject_sex is a one-hot vector indicating the sex of the subject animal (0 = female, 1 = male) and subject is a categorical variable indicating the subject animal. To avoid biasing the regression fit by the few cells with extremely low firing rates (indicated by circles in Figure S4A), we removed neurons with a more than 32-fold increase/decrease from the fit. The (llsubject) is a constant error term per subject animal, which we add due to the dependence introduced by the fact that we have unequal numbers of neurons recorded from the different subject animals^153^.

### Spiking neural network model

We simulated the activity patterns of ~1 mm^2^ of sensory cortex as 77,169 leaky integrate-and-fire (LIF) neurons with biologically realistic cell-type specific connectivity. The model was developed by ref. ^56^, has been validated several times^57–59^, and simulated spontaneous activity patterns, transmission delays and cell-type specific firing rates are in agreement with in vivo recordings in awake animals^56–57^. In our simulations, we modified the implementation of the model by ref. ^59^ and ran our simulations in the ‘Brian 2’ spiking neural network simulator for python^154^. The simulated neurons are distributed over eight populations, corresponding to excitatory and inhibitory neurons in layer 2/3, 4, 5 and 6. In addition, a population of thalamic neurons provide ‘touch’ input. The numbers of modeled neurons are given in Methods Table 1.

**Methods Table 1:**
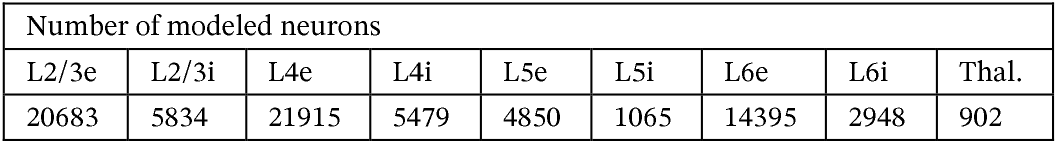
Number of modeled neurons.

The neural populations are connected with excitatory and inhibitory synapses. For every possible pre- to postsynaptic population, the model assigns K synapses

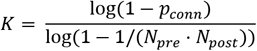

where *N*_*pre*_ and *N*_*post*_ are the total number of neurons in the populations (given in Methods Table 1) and p_conn_ is the connection probability (given in Methods Table 2). Once the total number of synapses is calculated, all the synapses are randomly assigned between neurons in the pre- and postsynaptic populations (pairs of neurons are drawn from a uniform distribution with replacement and *K* ≫ *N*, so there are multiple synapses per neuron).

**Methods Table 2:** connection probabilities

| | | Connection probability ($p_{\text{conn}}$) from... | | | | | | | | |
| --- | --- | --- | --- | --- | --- | --- | --- | --- | --- | --- |
|  |  | L2/3e | L2/3i | L4e | L4i | L5e | L5i | L6e | L6i | Thal. |
| Target | L2/3e | 0.10<br>1 | 0.1<br>69 | 0.0<br>44 | 0.0<br>82 | 0.03<br>2 | 0.0<br>0 | 0.0<br>08 | 0.0<br>0 | 0.0<br>0 |
|  | L2/3i | 0.13<br>5 | 0.1<br>37 | 0.0<br>32 | 0.0<br>52 | 0.30<br>75 | 0.0<br>0 | 0.0<br>04 | 0.0<br>0 | 0.0<br>0 |
|  | L4e | 0.00<br>8 | 0.0<br>06 | 0.0<br>5 | 0.1<br>35 | 0.00<br>7 | 0.00<br>03 | 0.0<br>45 | 0.0<br>0 | 0.09<br>83 |
|  | L4i | 0.06<br>9 | 0.0<br>03 | 0.0<br>79 | 0.1<br>60 | 0.00<br>3 | 0.0<br>0 | 0.1<br>06 | 0.0<br>0 | 0.06<br>19 |
|  | L5e | 0.10<br>0 | 0.0<br>62 | 0.0<br>51 | 0.0<br>06 | 0.08<br>3 | 0.37<br>3 | 0.0<br>20 | 0.0<br>0 | 0.0<br>0 |
|  | L5i | 0.05<br>5 | 0.0<br>27 | 0.0<br>26 | 0.0<br>02 | 0.06<br>0 | 0.31<br>6 | 0.0<br>09 | 0.0<br>0 | 0.0<br>0 |
|  | L6e | 0.01<br>6 | 0.0<br>07 | 0.0<br>21 | 0.0<br>17 | 0.05<br>7 | 0.02<br>0 | 0.0<br>40 | 0.2<br>25 | 0.05<br>12 |
|  | L6i | 0.03<br>6 | 0.0<br>01 | 0.0<br>03 | 0.0<br>01 | 0.02<br>8 | 0.00<br>8 | 0.0<br>66 | 0.1<br>44 | 0.01<br>96 |

The membrane potential, *V*(*t*), of an LIF neuron is governed by the differential equation

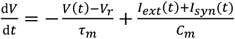

where *τ*_*m*_ = 10 ms is the membrane time constant, *C*_*m*_ = 250 pF is the membrane capacitance, *V*_*r*_ = −65 mV is the reset potential, *I*_*exp*_(*t*) is a background current and *I*_*syn*_ (*t*) is the synaptic current. When the membrane potential of a neuron reaches a threshold value, *V*_*th*_ = −50 mV, the neuron spikes and is reset and held at a reset voltage, *V*_*r*_ = −65 mV, for a refractory period of *τ*_*ref*_. = 2 ms. When a presynaptic neuron spikes, the postsynaptic neuron receives an increase in synaptic current of *w*_*e*_ (the synaptic weight) for excitatory neurons and *w*_*i*_, = −4*w*_*e*_ for inhibitory neurons. *w*_*e*_ is drawn from a normal distribution, *Normal*(*μ* = 87.8 pA, *σ* = 8.8 pA). The synaptic weight for connections from neurons in L4e to L23e is doubled. Synaptic currents are delivered with a delay *d_e_ = Normal(*μ* =* 1.5 ms, *σ* = 0.75 ms) for excitatory synapses and *d*_*i*_ = Normal(*μ* = 0.8 ms, *σ* = 0.4 ms) for inhibitory synapses. Synaptic current changes are modeled as exponentials with a time constant *τ*_*syn*_„ = 0.5 ms and and governed by the differential equation,

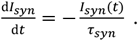

We initialized the simulation with an initial membrane potential drawn from *Normal*(*μ* = −58 mV, *σ* = 10 mV). The network is driven by delivering a layer-specific background current to all neurons, *I*_*ext*_(*t*) = *b* · 0.3512 pA, where b is a layer-specific background scaling factor. The background scaling factor, b, is given in Methods Table 3.

**Methods Table 3.** Background input.

| Background scaling factor (b) |  |  |  |  |  |  |  |
| --- | --- | --- | --- | --- | --- | --- | --- |
| L2/3e | L2/3i | L4e | L4i | L5e | L5i | L6e | L6i |
| 1600 | 1500 | 2100 | 1900 | 2000 | 1900 | 2900 | 2100 |

At baseline, the network does not receive input from the thalamic population (firing rate of thalamic neurons, *r*_*thal*_ = 0 Hz). Each simulated touch trial started with a baseline period pf 1000 ms, then we simulated ‘touch’ by increasing *r*_*thal*_ to 30 Hz for a period of 700 ms, and then we again simulated a baseline period of 300 ms (see Figure 5b for two example trials). In eveiy other trial, we simulated a change in the network, stemming from social-touch-mediated release of oxytocin during touch. In order to simulate a depolarization-induced increase in firing rate of interneurons during touch, we increased the reset potential, *V*_*r*_, of the intemeuron populations by [0 mV, + 0.5 mV, +1.0 mV], In order to simulate an increase in input resistance of excitatory neurons during, we multiplied the membrane time constant, *τ*_*m*_, with [1.0,1.1,1.2], only during touch. We scale the membrane time constant *τ*_*m*_, since *τ*_*m*_ = *R*_*m*_ · *C*_*m*_, where *R*_*m*_ is the membrane resistance and *C*_*m*_ is the membrane capacitance. All simulations were done using the ‘Brian 2’ exact integration method for linear equations, with 0.1 ms time steps. For eveiy configuration, we performed 20 non-modulated and 20 modulated trials (interleaved) and calculated *β*_*touch*_ = log(*r*_*touch*_/*r*_*base*_) for 2000 neurons chosen randomly across all populations. We estimated the rates *r*_*base*_ and *r*_*touch*_ by counting spikes in a 700-ms window before and after the simulated touch onset, respectively. We removed extremely low-firing neurons (neurons with less than three spikes across trials). In order to characterize the network response pattern, we fit three models to the modulated and non-modulated responses: a ‘Bias model’ with only a bias and the slope fixed at unity; a ‘Potentiation model' with no bias, but a free slope parameter; and a ‘Full model’ with both a bias and a potentiation of responses:

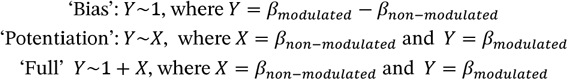

The models were fit using the ‘statsmodels’ python package. For all three models, we calculated the Bayesian information criterion and used this to select the model, which best characterizes the data (indicated by a red dot in the small inserted plots in Figure 5c).

### Identification of putative inhibitory/excitatory neurons by extracellular spike shape

As a low-level check of our network simulations, we compared the touch responses of putatively excitatory and inhibitory neurons in somatosensory cortex (based on their extracellular spike shape) with the response of simulated excitatory and inhibitory neurons in the model. In some brain regions, for example the hippocampus, separation of cell type based on extracellular spike shape appears to work well and to reliably identify thin-spiked neurons which suppress simultaneously recorded neurons with short latency^155,156^. However, in other brain regions the use is more controversial^157^. For example, motor cortical pyramidal projection neurons have extremely thin spikes^158^ and some cortical interneurons have very wide spikes ^159,160^. Further, since the spike width and shape depend strongly on the relative location of the electrode and the cell^161^, spike shapes recorded with tetrodes in the agranular rat frontal cortex (with the lack of cytoarchitectonic stereotypy) are likely less reliable than spike shapes recorded in the hippocampus ^l48,162^. Following our previous report on ST^29^, we used two features of the spike shape: the spike width (at half maximum) and the ‘post-positivity’ (the integral of the spike waveform between 0.375 ms and 0.75 ms after the spike peak, normalized by peak voltage). These are slightly different features from our previous report, focusing on Al^33^. For feature calculations, we oversampled the average spike at twice the sampling rate with cubic spline interpolation. For clustering, we standardized both features (by z-scoring) and fitted a Gaussian mixture distribution with two components (‘fitgmdist’ in Matlab, with a regularization of 0.1) and clustered the data based on the fitted components. For SI (Supplementary Figure 7b-c) and Al (Supplementary Figure 7e-g), this approach was feasible: the 2d-distribution of features was bimodal, and we could fit a two-component Gaussian mixture model. As we have previously reported^34^ - and in line with the motor-cortex specific caveats outlined above - we did not see a bimodal 2d-distribution of features in the vibrissa motor cortex data, and we could not fit two well-separated Gaussian components. Thus, here we simply fitted a single 2d Gaussian, and ‘cut’ the 2d-distribution through the mean, along the shortest axis (Supplementary Figure 7h-j). We excluded seven neurons from this analysis because the spike shapes were either not properly stored in our database (six Al neurons) or where the spike shape was contaminated with noise because a nearby neuron often spiked close by in time (one VMC neuron). As a general word of caution, we would like to highlight that in our previous analysis of the tetrode recordings from somatosensory cortex, separating neurons into putative inhibitoiy and excitatory neurons by extracellular spike shape suggested that social touch responses of regular-spilcing neurons in somatosensory cortex changed across the estrus cycle ^29^. However, in a follow-up study usingjuxtacellular recordings (which capture the spike shape with much higher fidelity than tetrodes), we were unable to reproduce that finding^31^. Even in brain regions where spike shape features appear bimodal, such as SI, they are only a weak proxy for cellular identity, at least in cortex^157^.

### Data and code availability

Matlab code and a table including the response magnitudes from the fitted ‘sex-touch’ models, the sex and identity of the subject animal (i.e. all the data needed to reproduce Figure 4 and reproduce the statistical modeling) is available as supplementary material. All python code required to run the simulations shown in Figure 5, and the python code to analyze the simulated data and generate the figure panels are available as supplementary material. The full raw data from four example neurons (the neurons shown in Figure 3), as well as the Matlab code used to fit and analyze the statistical models and generate plots in Figure 3 is available as supplementary material. The full raw dataset is available upon request to the corresponding author.

## Acknowledgements

We thank Gabriel Curio for valuable discussions. We thank Sara Helgheim Tawfiq for behavior drawings. This work was supported by The Novo Nordisk Foundation (C.L.E), Humboldt Universität zu Berlin within the Excellence Initiative of the states and the federal government (C.L.E), BCCN Berlin (German Federal Ministiy of Education and Research BMBF, Förderkennzeichen 01GQ1001A) (M.B.), Humboldt-Universität zu Berlin (M.B), NeuroCure (M.B.), and the Gottfried Wilhelm Leibniz Prize of the DFG (M.B.).

## Author contributions

C.L.E., E.B. and R.P.R. provided clustered unit data and behavior data from previous studies. C.L.E. performed additional tetrode experiments. C.L.E. designed and performed analysis and statistical modeling, designed and performed spiking neural network simulations and prepared the figures. M.B. supervised the study. C.L.E. wrote the first version of the manuscript. All authors contributed to writing the paper.

## Competing interest statement

The authors declare no competing interests.

---

**Supplementary Figure 1.**
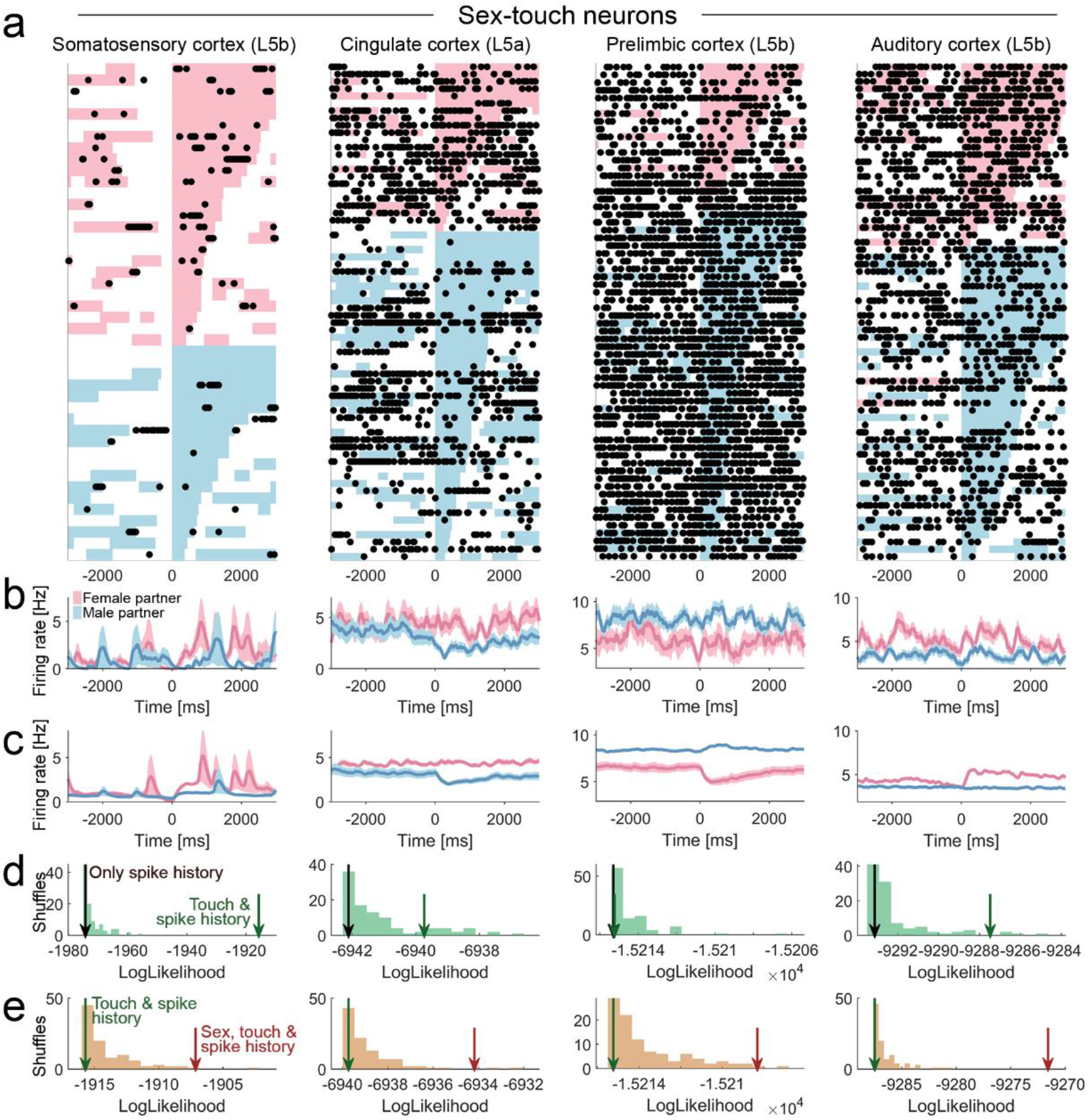
More example sex-touch neurons from the other brain areas. (a) Raster plot of example sex-touch neurons for the brain areas not shown in Figure 2 (an S1 L5b neuron, an ACC L5a neuron, a PrL L5b neurons and an A1 L5b neuron). Raster plots show spike times (black dots) aligned to the first whisker-to-whisker touch in each social touch episode. Social touch episodes are sorted by partner sex (female: pink, male: blue) and by duration (indicated by length of colored bar). (b) Peri-stimulus time histograms of the example neurons shown in (a), separated by partner sex. Black line indicates mean firing rate, smoothed with a Gaussian kernel (*σ* = 100 ms), shaded area indicates s.e.m, pink/blue color indicates female/male partner animals. (c) Peri-stimulus time histograms of the example neurons shown in (a), calculated from the fitted regression model (plot conventions as in (b)). (d) Estimating touch-modulation: Log-likelihood values of models fitted to the neurons in (a). The log-likelihood of models depending on touch is indicated by a green arrow, the log-likelihood of models without touch is indicated by a grey arrow and the log-likelihood distribution of shuffled touch-models is indicated by green bars. Some sex-touch neurons would not be significant, if the partner sex was not considered in the model. For example, the ACC L5a neuron is suppressed by males, but shows (almost) no response with females, so the green arrow is not in the 0.05 fraction of the shuffled distribution if all touches are pooled, without parsing out the partner sex. (e) Estimating sex-touch-modulation: Log-likelihood values of models fitted to the neurons in (a). The log-likelihood of models depending on partner sex and touch is indicated by a brown arrow, the log-likelihood of model without sex is indicated by a green arrow and the log-likelihood distribution of shuffled sex-touch-models is indicated by brown bars. All neurons are significant at p < 0.05 (brown arrow outside the shuffled distribution).

**Supplementary Figure 2.**
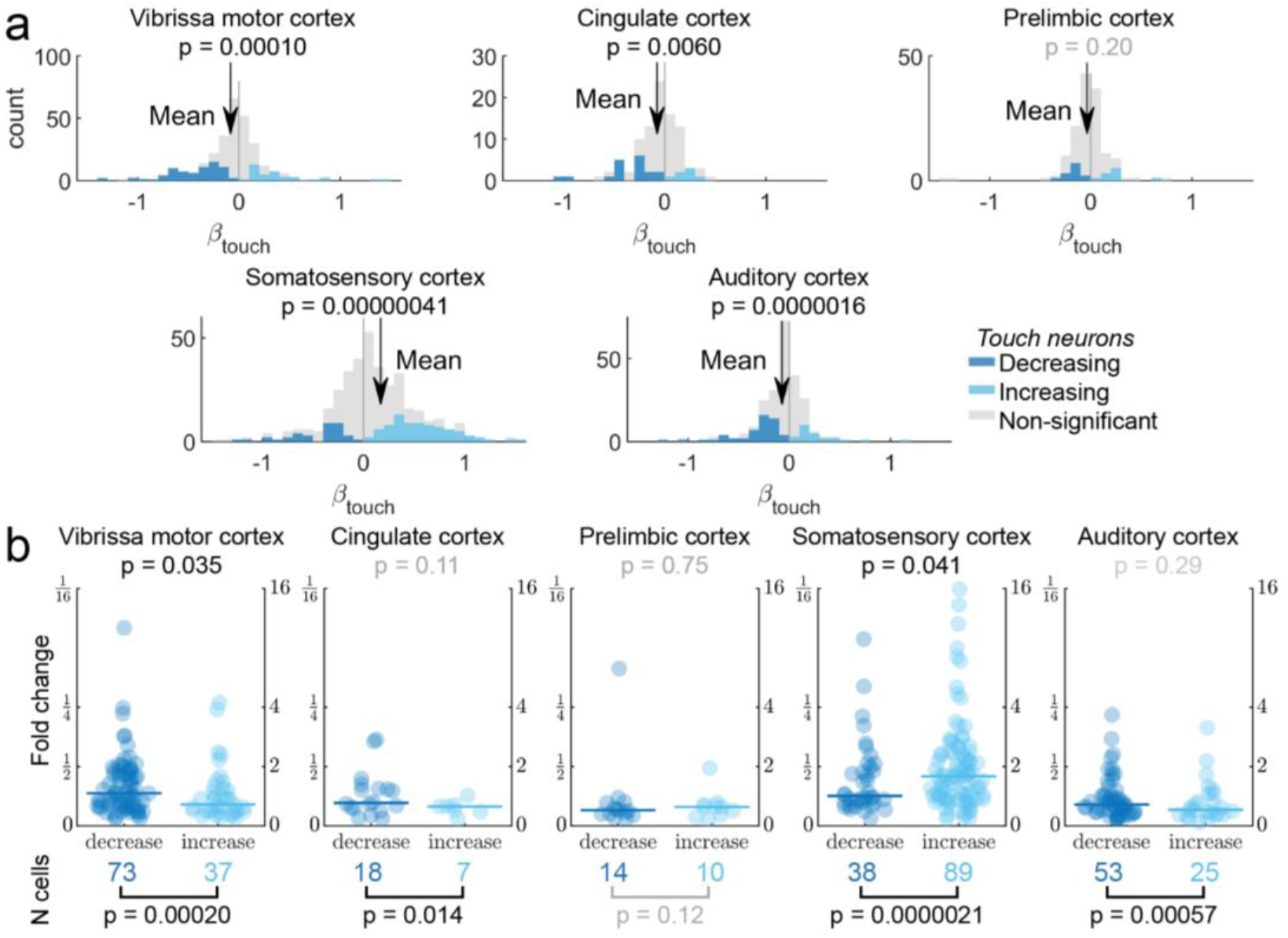
Prototypical responses during social touch vary by cortical area. (a) Distribution of fitted *β*_*touch*_ across cortical areas (axis clipped at +/−1.6, all data used for calculations, all data plotted in S2A-B). Colored bars indicate significantly increased neurons (light blue), significantly decreased neurons (dark blue) and non-significant neurons (grey). Black arrow indicates mean *β*_*touch*_, p-value indicates Wilcoxon signed rank test. As a population, S1 neurons increased in firing rate during social touch (mean *β*_*touch*_ = 0.17, p = 0.00000041, N = 384, Wilcoxon signed-rank test). VMC, ACC and A1 neurons decreased (VMC/ACC/AC: mean *β*_*touch*_ = −0.084/-0.077/-0.073, p = 0.00010/0.0060/0.0000016, N = 296/95/239, Wilcoxon signed-rank test) and PrL neurons did not show any modulation at the population level (mean *β*_*touch*_ = −0.036, p = 0.20, N = 142, Wilcoxon signed-rank test). (b) *Top:* Fold change in firing rate by social touch for significantly increased (light blue) and decreased neurons (dark blue). Horizontal lines indicate medians, p-values indicate Mann-Whitney U-test. *Below:* Number of neurons, which are significantly increased (light blue) and decreased (dark blue) by social touch. P-values indicate binomial test. In S1, more neurons were significantly increased by touch (decreasing v. increasing neurons, 38 v. 89, p = 0.0000021, binomial test) and the significantly increasing neurons were the most strongly modulated neurons (decreasing v. increasing neurons, median |log2(ratio)| = 0.51 v. 0.84, p = 0.041, Mann-Whitney U test). In VMC, more neurons were decreasing (73 v. 37 neurons, p = 0.00020, binomial test) and the decreasing neurons were most strongly modulated In VMC (median |log2(ratio)| = 0.55 v. 0.36, p = 0.035, Mann-Whitney U test). ACC and A1 both had more significantly decreasing neurons (ACC: 18 v. 7 neurons, p = 0.014, AC: 53 v. 25, p = 0.00057, binomial test), but no significant difference in the modulation strength. (ACC: median |log2(ratio)| = 0.39 v. 0.33, p = 0.11, A1: median |log2(ratio)| = 0.36 v. 0.27, p = 0.29, Mann-Whitney U-test). PrL had a similar number of increasing and decreasing neurons and no differences in modulation strength (14 v. 10 neurons, p = 0.12, binomial test, median |log2(ratio)| = 0.26 v. 0.32, p = 0.75, Mann-Whitney U test).

**Supplementary Figure 3.**
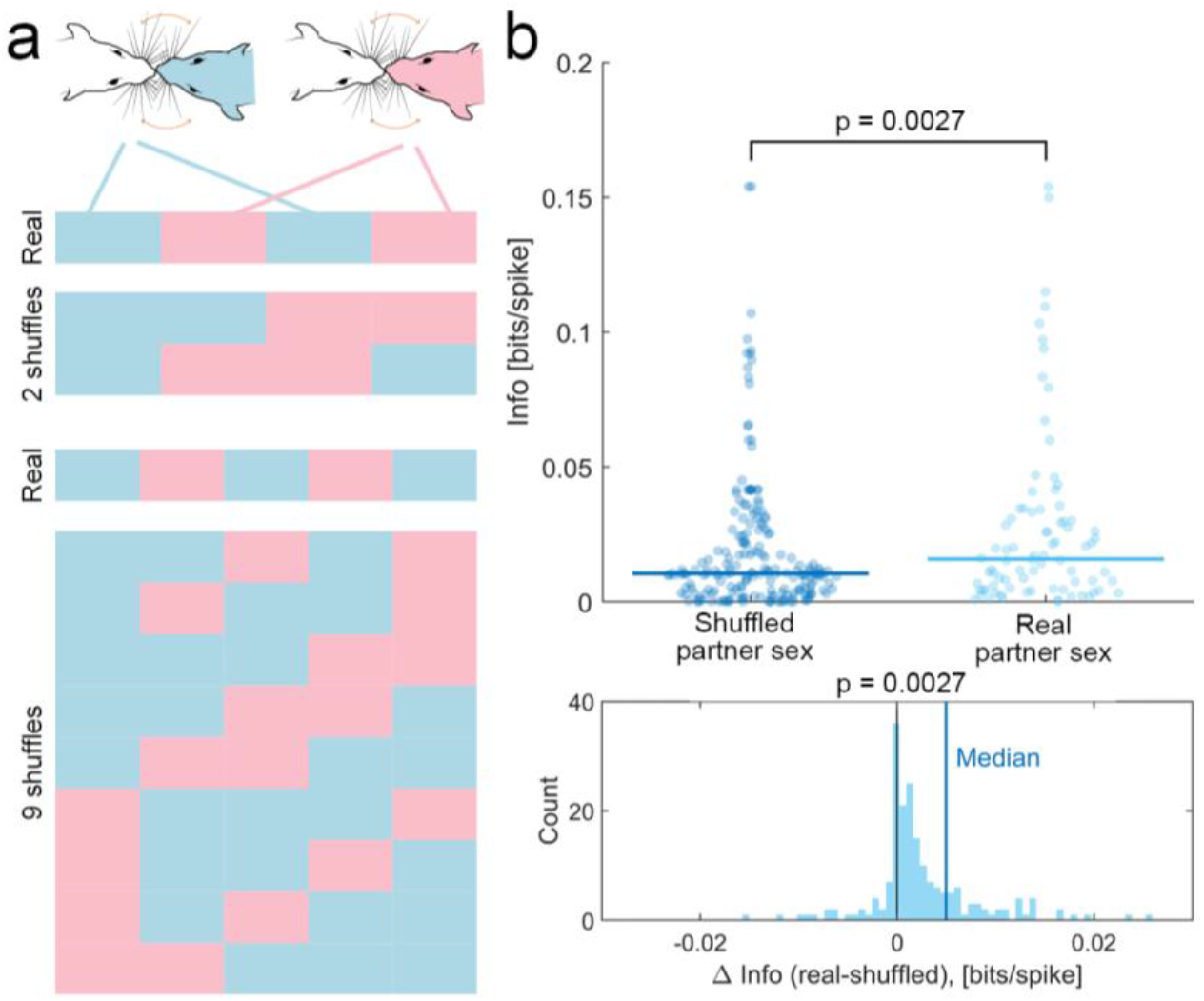
Partner sex patterns responses more than individual identity. If neurons did not encode the sex of the stimulus animals at all, but had individual-specific responses, we would expect to sometimes identify artefactual ‘sex-touch’ neurons, simply because we are comparing two groups of animals with individual responses. Since we only presented few partner animals, it is difficult to ask (at the single-cell level) if responses of these neurons are individual-specific or sex-specific. One way to ask if ostensibly partner-sex-dependent response differences are indeed driven by the sex of the partner animal is to ask if grouping the responses to partner animals by their real sex is more informative about spike rates, than grouping the responses to partner animals by a shuffled sex. We frame the analysis in terms of mutual information between spike rates, because this allows us to be agnostic about the direction of modulation (increases/decreases). First, we used a shuffling procedure to identify neurons, most likely to carry individual-specific information (see Methods). The number of possible shuffles depends on the number of partner animals in the particular recording session (typically 2 males and 2 females, panel a). To overcome this imbalance in the data, we used a mixed-effects modeling approach^9^ (panel b). We found that putatively individual-specific neurons were significantly more informative about sex than all other possible partitions of the data (mean Δlnfo = 0.007 bits/spike, p = 0.0027, mixed-effects model, panel b). This analysis does not exclude the possibility that some neurons with individual-specific responses might still be present in these brain areas. However, it shows that the sex of the partner animal is indeed a major determinant of the firing patterns, and that partner-sex-specific modulation is not an artifact better explained by individual-specific effects. (a) Number of possible shuffles grows with number of interaction partners. If an animal interacted with two male and two female conspecifics, we can only generate two possible shuffled assignments of the partner sex (top). If the animal interacted with three males and two females, we can generate nine possible shuffled assignments of the partner sex (bottom). (b) Top: Distribution of information per spike calculated using real (light blue) and shuffled (dark blue) assignments of partner sex. Below: Distribution of difference between real and shuffled information per spike (p-value indicates mixed-effects model, see Methods).

**Supplementary Figure 4.**
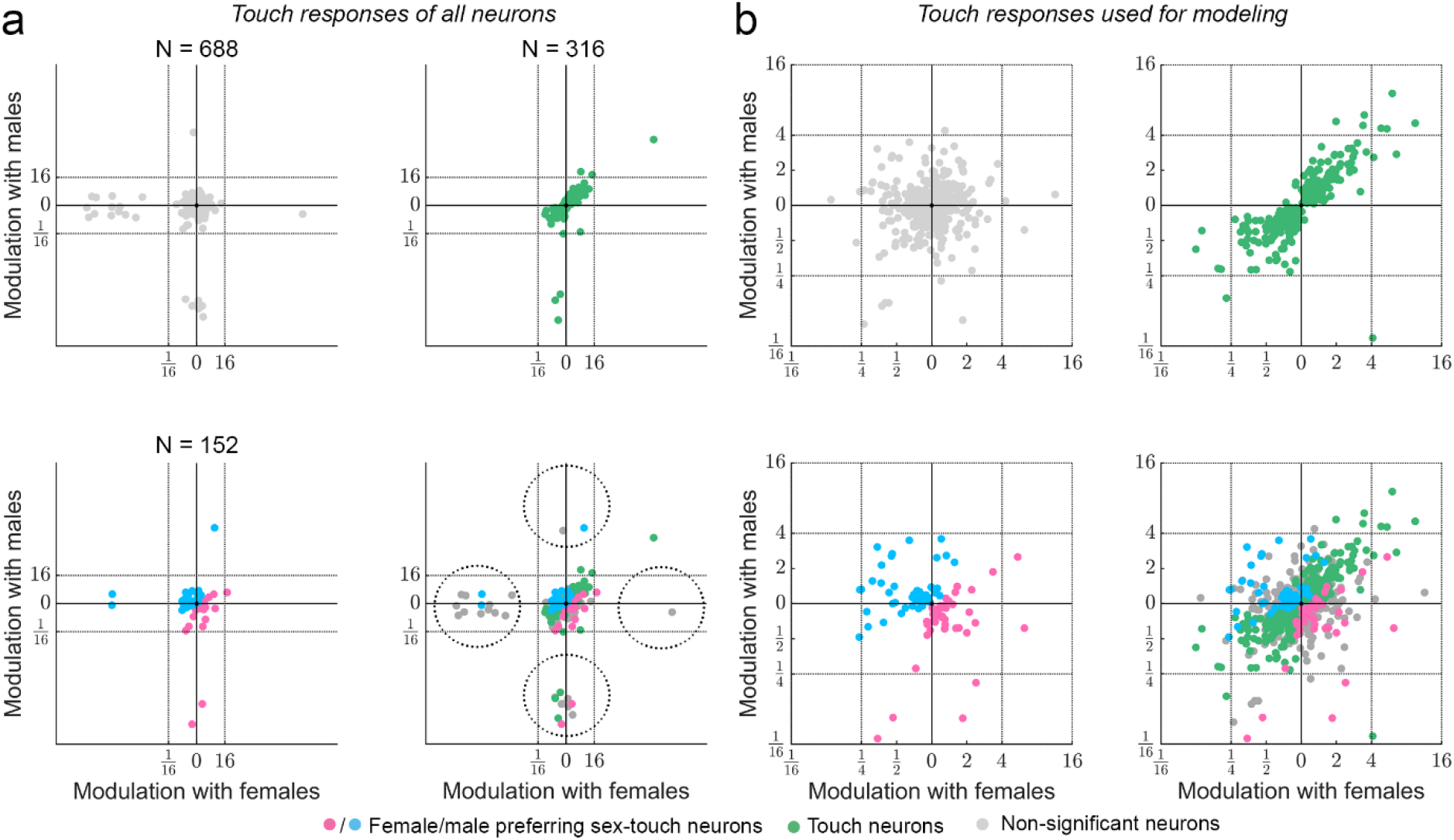
Population response patterns across all areas. (a) Modulation of activity (in fold change) of all neurons during social touch with male and female conspecifics. Touch neurons are indicated by green dots, female/male preferring sex-touch neurons are indicated by pink/blue dots and non-significant neurons are indicated by grey dots. Since the statistical modeling of the spike train models all modulation as ratios, a few neurons with either very low baseline firing rates, or which were essentially silenced during touch will be fitted to very high/low values of modulation (indicated by dotted circles). (b) Same plots as (a), but zoomed in to only show neurons between 16-fold increase and 16-fold decreases in firing rate (the vast majority of neurons). In order not to skew the GLM models by the extreme outliers, we only used neurons with less than 32fold modulation in the GLM models. Touch neurons are indicated by green dots, female/male preferring sex-touch neurons are indicated by pink/blue dots and non-significant neurons are indicated by grey dots.

**Supplementary Figure 5.**
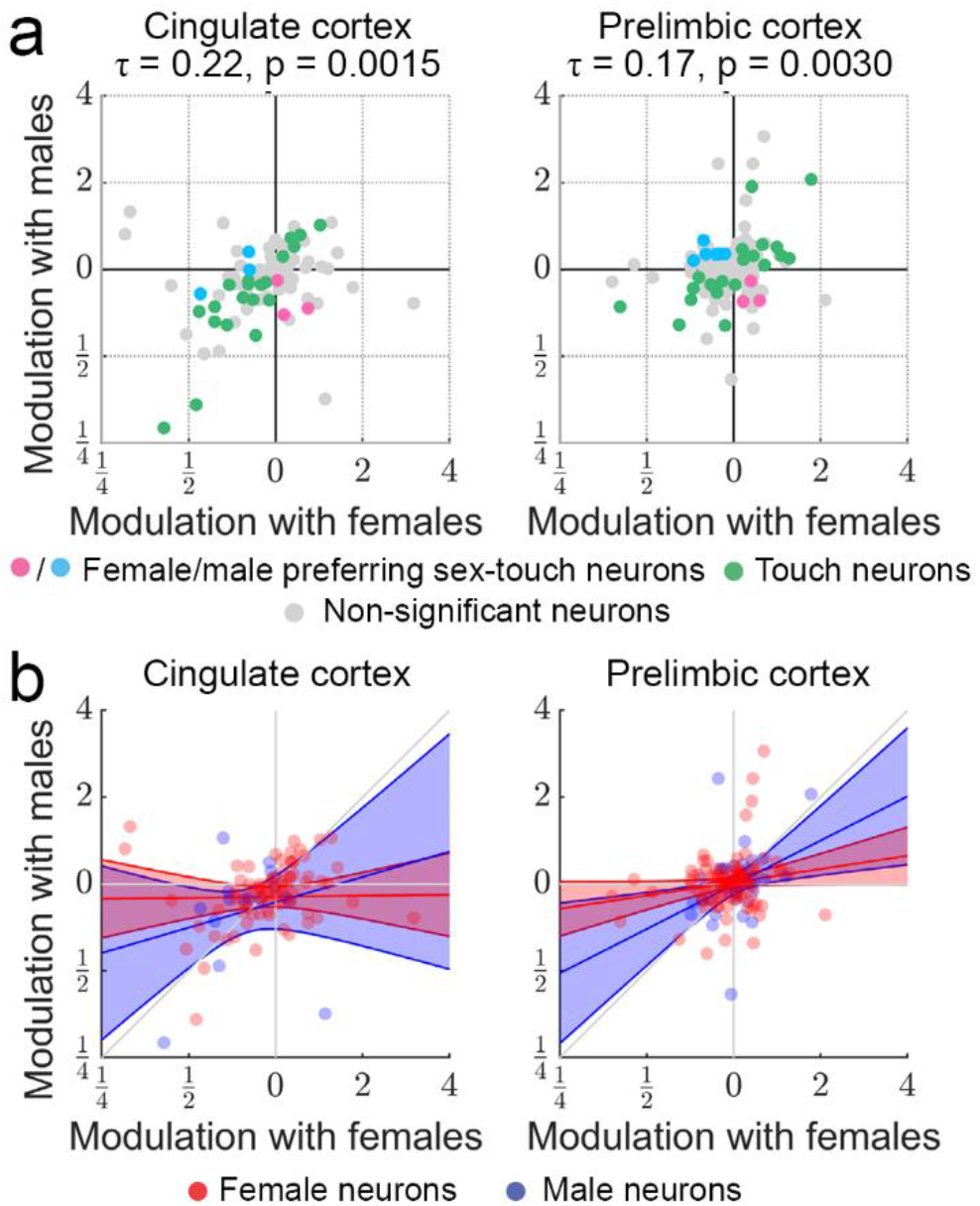
Population response patters during social facial touch for cingulate and prelimbic cortex. (a) Same plot as Figure 4c, but showing data from ACC and PrL. Modulation of activity (in fold change) during social touch with male and female conspecifics is highly correlated. Touch neurons are indicated by green dots, female/male preferring sex-touch neurons are indicated by pink/blue dots and non-significant neurons are indicated by grey dots, Kendall’s *τ* and p-value above. (b) Same plot as Figure 4d, but showing data from ACC and PrL. Although not significant, the maximal-likelihood fit also estimated *β*_*subject_sex*_ to be less than unity for both ACC and PrL. This pattern is in line with the pattern in S1, VMC and AC (Figure 4d). Red/blue dots indicate neurons recorded in female/male subject animals, red/blue lines indicate maximum-likelihood fit of regressing modulation with males as a function of modulation with females, for female/male subjects (see Methods and Suppl. Note 2 for model specification), shaded area indicates 95% C.I.

**Supplementary Figure 6.**
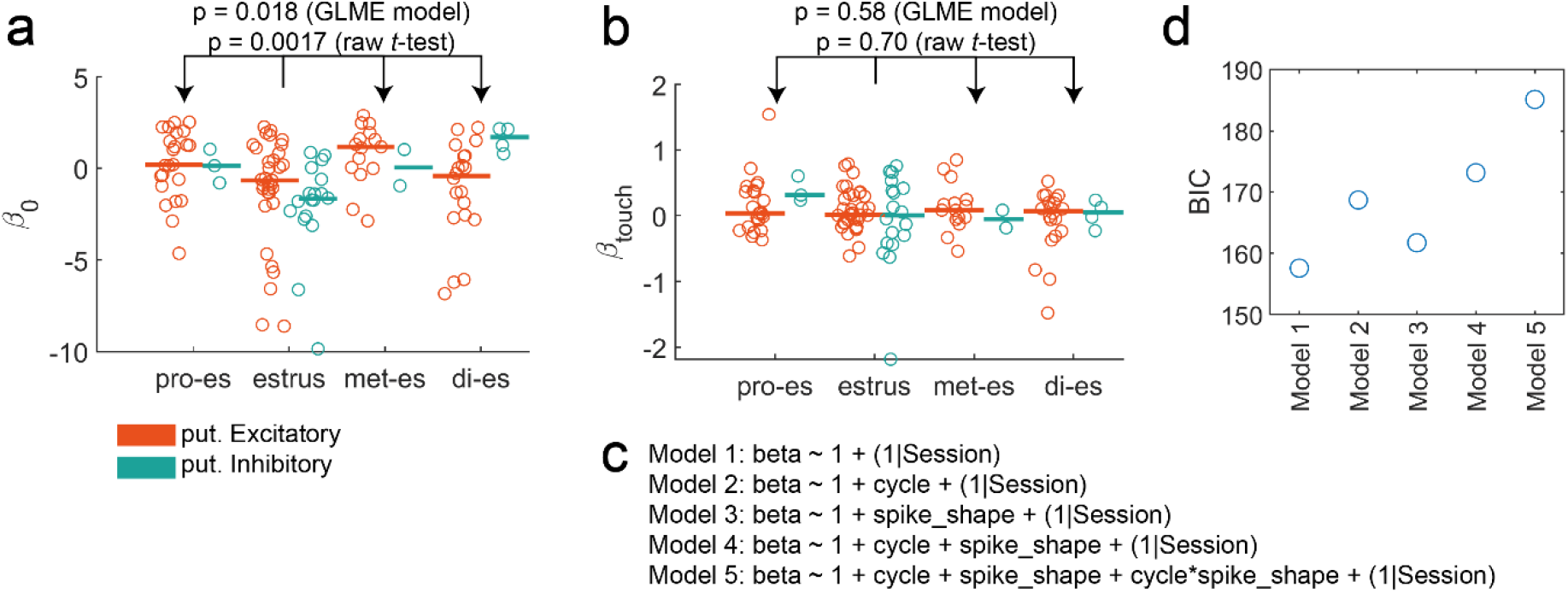
Responses to social facial touch in somatosensory cortex do not depend on the estrus state. (a) In a subset of the S1 data from female subject animals, we had access to the estrus state at the day of the experiment. In agreement with ref. ^4^ (same data) and ref. ^10^, we found that firing rates (quantified here as *β*_0_, so baseline rate *r*_*base*_ = exp(*β*_0_)) were significantly lower during estrus than non-estrus. We did two tests: a “raw” t-test where we pooled all non-estrus states (like in ref. ^4^) and tested against the estrus state cells (p = 0.0017), and a more sophisticated model, where we treat all non-estrus days (pro-estrus, met-estrus and di-estrus) as independent categorical variables, control for unequal number of neurons recorded on the same experimental session, fit a GLME regression and do the full ANOVA (p = 0.018). Both were significant. For plotting, we have split putative excitatory (orange) and inhibitory (teal) neurons, vertical lines indicate medians. (b) Same analysis as in (a), but here we analyze the responses to social touch (quantified as *β*_*touch*_). We did not find any differences in response magnitude across the estrus cycle (neither using a pooled t-test or the GLME model: p ≫ 0.05, same finding as ref. ^10^). (c) We tried a variety of GLME models to see if responses depended on estrus state (‘cycle’, 4 levels) or putative inh./ex. neuron type (‘spike_shape’, 2 levels, shown in Figure S7 and S8), or both (interactions), but none of these models had significant effects or did better than a constant model (models compare by Bayesian information criterion, model specification in Wilkinson notation). (d) Bayesian information criteria (BIC) for the models shown in (c). The constant model (‘Model 1’) has the lowest BIC.

**Supplementary Figure 7.**
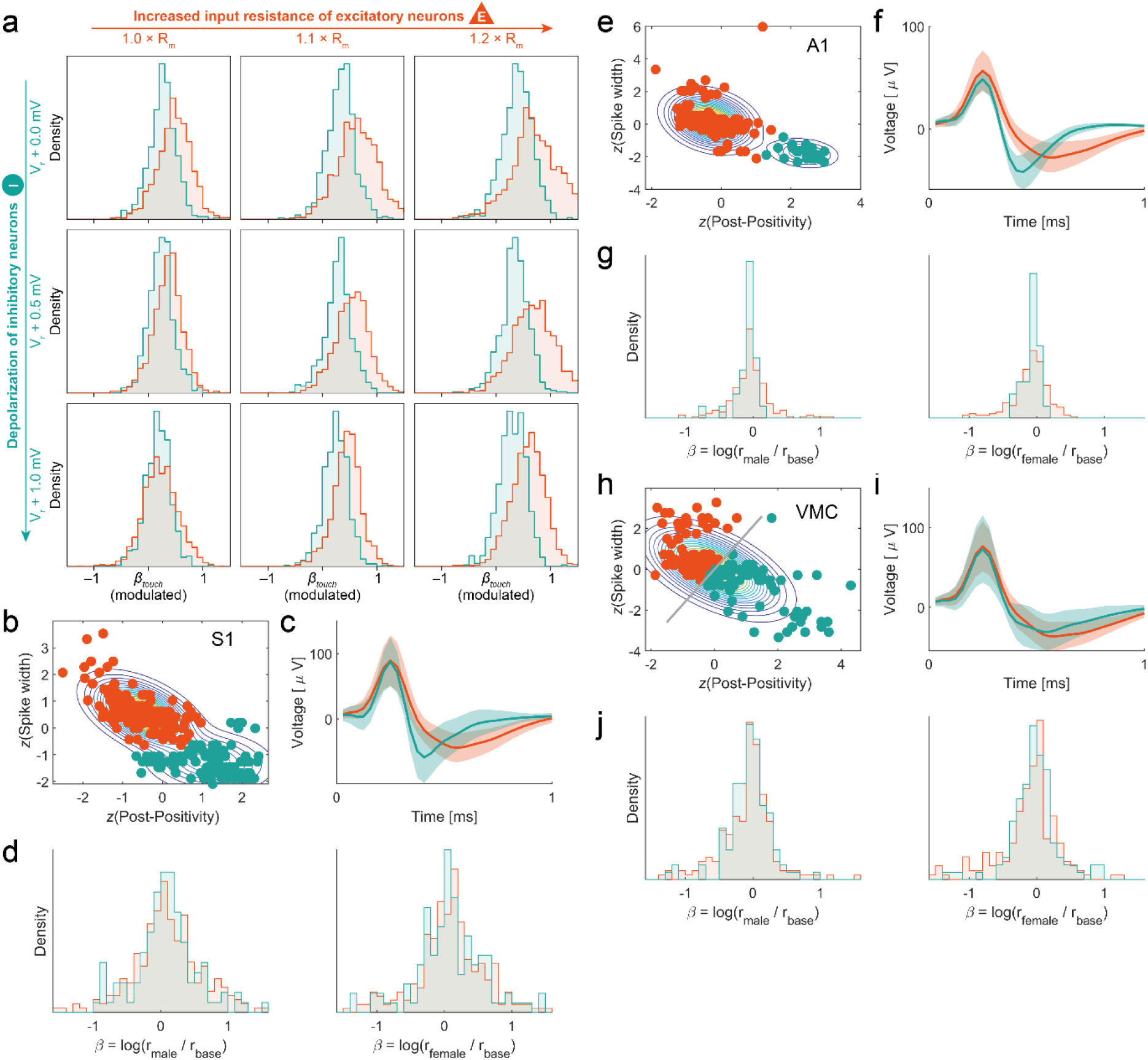
Putatively inhibitory and excitatory neurons respond in the same direction and overlap, consistent with simulated responses to ‘touch’ (thalamic input). (a) Density of responses during touch of simulated excitatory (orange) and inhibitory (teal) neurons are in the same direction an overlap (more and more strongly, going down along first column, same plotting conventions as Figure 5c). (b) We separated putatively excitatory (orange) and inhibitory (teal) neurons recorded in somatosensory cortex by two features: the spike width (at half maximum) and the ‘post-positivity’ (the integral of the spike waveform between 0.375 ms and 0.75 ms after the spike peak, normalized by peak voltage). We assigned the neurons two clutsers by z-scoring the features, fitting a two-component Gaussian distribution and assigning the neurons by the component yielding the highest posterior probability (see Methods). (c) Mean spike shape of putatively excitatory (orange) and inhibitory (teal) neurons recorded in somatosensory cortex (shaded area indicates standard deviation).(d) Density of responses during social touch to male and female conspecifics of putatively excitatory (orange) and inhibitory (teal) neurons recorded in somatosensory cortex. (e-g) Same as (b-d), but for neurons recorded in auditory cortex. (h-j) Same as (b-d), but for neurons recorded in vibrissa motor cortex. Note in panel h, the spike widths and post-positivity did not form a bimodal distribution, so we just fitted a single multivariate Gaussian and ‘cut’ that gaussian in two along the shortest axis (grey line, please see Methods).

## Supplementary Note 1: A brief introduction to β-coefficients as a metric of firing rate changes

In this study, we want to investigate how social touch and social context impact the firing patterns of cortical networks. Thus, we need a metric which allows us to quantify changes in firing rate of both single neurons and populations of neurons. In this short note, we explain why using regression coefficients (“*β*-coefficients”) as metric is a natural choice. We also compare the *β*-coefficients to other commonly used metrics of firing rate change.

### Not all metrics are suitable for quantifying patterns in population activity

Let us first consider a ‘classic’ peri-stimulus time histogram (PSTH). For a cartoon neuron, a PSTH aligned to the beginning of social touch might look like this:

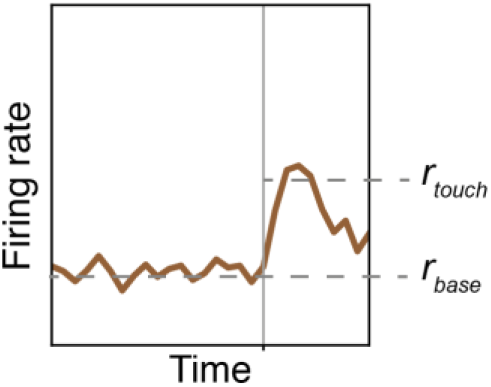

Which metric should we use to quantify how much the firing rate changes during touch? One possibility would be to estimate a baseline firing rate (*r*_*base*_) and a firing rate during touch (*r*_*touch*_) and simply measure the change in firing rates: *Δr* = *r*_*touch*_ − *r*_*base*_. This metric does not take the baseline firing rate into account, and it is thus difficult to compare across neurons. For example, for these two cartoon neurons, *Δr* is the same, but it seems right to say that the neuron on the left is more “strongly modulated” than the neuron on the right:

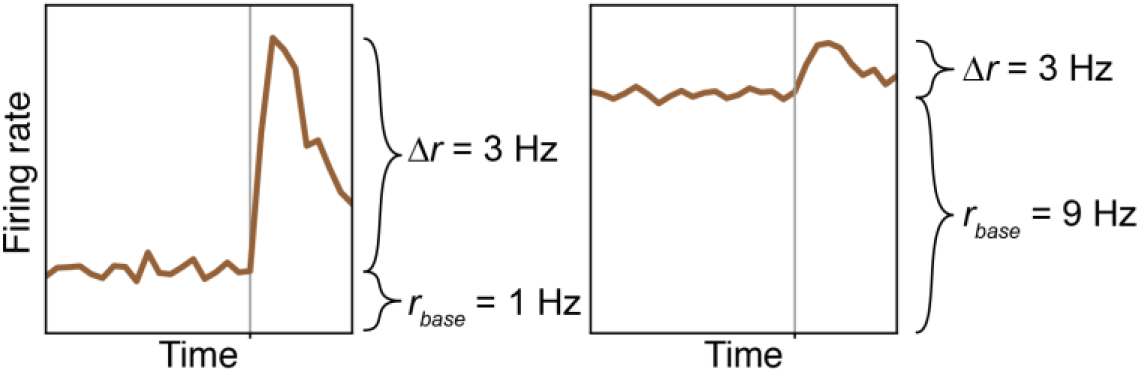

The z-score is another commonly used metric of firing rate changes. In a PSTH-based analysis, the z-score measures how much the firing rate during touch increases above the baseline firing rate, measured in multiples of the standard deviation of the baseline firing rate:

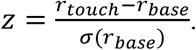

Like *Δr* the z-score does not normalize by the baseline firing rate and it is thus also difficult to compare *z*-scores across neurons. Moreover, since z-scores are normalized by the standard deviation of the baseline firing rate, z-scores of a PSTH depends on the number of trials. For example, even though these two cartoon neurons respond equally to touch, the left neuron will have a small z-score (there are few trials, so the PSTH is noisy and has a high standard deviation), whereas the right neuron will have a large *z*-score (there are many trials, so the baseline firing rate is very smooth and flat):

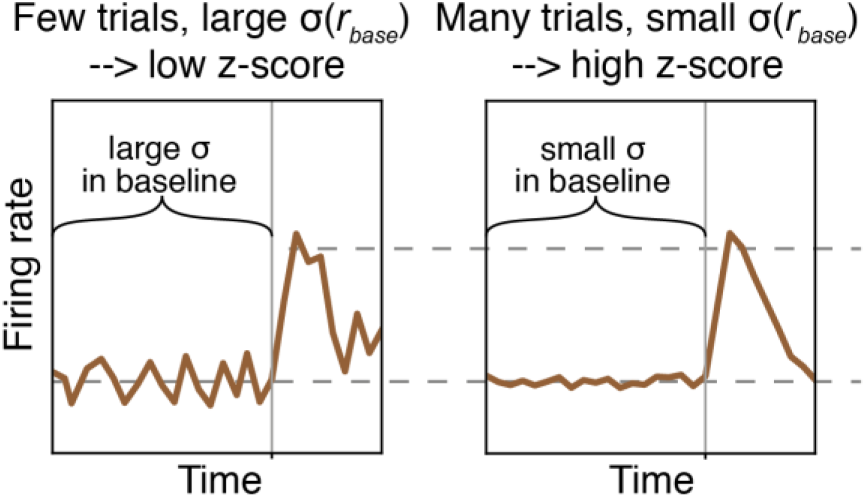

### We need a metric which takes baseline firing rates into account

Clearly, both *Δr* and z-scores are not good metrics if we want quantify the modulation of a whole population of neurons. We need a metric that normalizes the change in firing rate by the baseline firing rate. Let us consider three options:

1. One option is to quantify the modulation a neuron using the metric 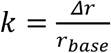. The interpretation of this metric is simply the magnitude of change measured in units of the baseline rate. Firing rates cannot be negative (*r* ≤ 0Hz), so this metric takes values in the interval [−1, ∞[. For neurons with decreasing firing rates, *k* will be between −1 and 0, whereas for neurons with increasing firing rates, *k* can be arbitrarily large.
2. A second option would be to quantify modulation using the metric 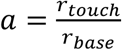. The interpretation of this metric is as the ratio of firing rate during touch and baseline. This metric takes values in the interval [0, ∞[. For neurons with decreasing firing rates, *a* will be between 0 and 1, whereas for neurons with increasing firing rates, *a* can be arbitrarily large.
3. A third option is to wrap the ratio of firing rates inside some function. Any function could be used in our metric, as long as that function is injective (every *f(a)* is unique for all possible values of *a*). For example, we can choose the logarithmic function, and define our metric as 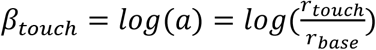. Because we used the logarithm, the interpretation of the metric *β*_*touch*_ is straightforward: *β*_*touch*_ is a ‘fold change’. This metric takes values in the interval] − ∞, ∞[^1^. In contrast to *k* and *a* (where increases in firing rates lead to arbitrarily large values of the metric, but decreases in firing rates were assigned values between −1 and 0, or 0 and 1), *β*_*touch*_ is symmetric around 0. Doubling the firing rate during touch (a = 2) and halving the firing rate during touch (*a* =½) yield *β*_*touch*_ of equal magnitude, because both are a “two-fold” change. Since we used the natural logarithm, *β*_*touch*_ = ± 1 corresponds to an e-fold increase and an e-fold decrease, respectively:

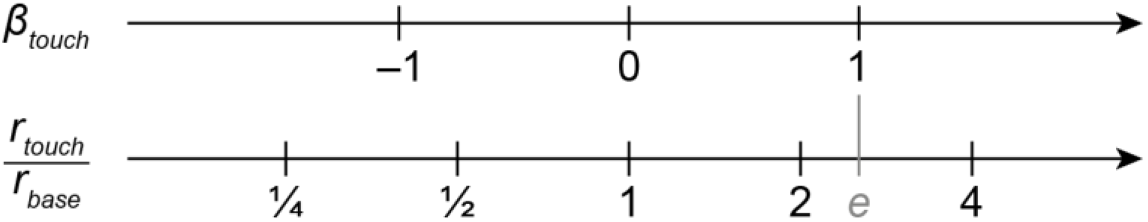

### What are the reasons to prefer one metric over the other?

We have introduced three metrics, which quantify how the firing rates of single neurons change during episodes of social touch:

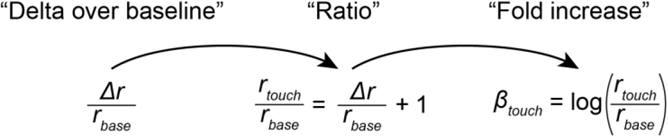

These metrics all measure the same thing and one can be calculated from the other, so what is the reason to prefer one over the others? The reason is the shape of their distribution across the network. In this study, we want to quantify patterns in the activity in large populations of neurons, and ask how these patterns depend on sex of the partner animal and sex of the subject animal. Such questions require analysis of the variance of our metric. Generalized linear mixed-effect modeling (GLME models) is a powerful statistical method that allows us to perform such analysis of variance (ANOVA), but such regression methods are only valid if we can model the error distributions. Specifically, if the errors follow a Gaussian distribution, or at least an approximately Gaussian distribution, we can use GLME modeling to analyze how touch responses depend on partner sex and subject sex, while simultaneously controlling for the fact that we have an unequal number of neurons from the experimental subject animals^9^.

So which metric is a natural choice for cortical firing rates? That is an empirical question. A first clue is provided by the fact that baseline firing rates across neural populations are usually log-normally distributed^13^. This is also the case in our dataset. The distribution of firing rates in our data is heavily skewed towards low firing rates with a heavy tail of few, high-firing neurons. If we plot the distributions of firing rates on a logarithmic axis, the distribution is approximately Gaussian (best Gaussian fit is plotted on top):

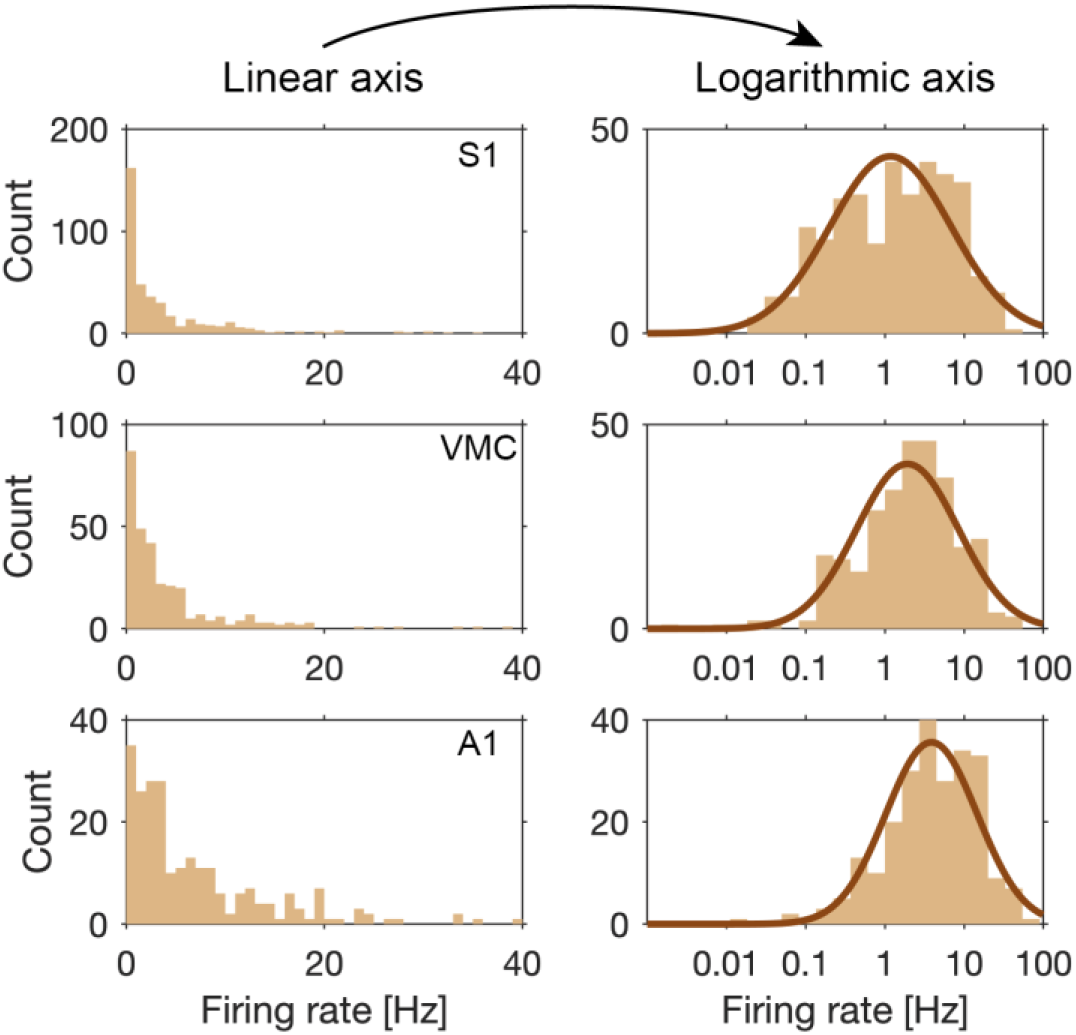

Since firing rates are log-normally distributed, it seems likely that changes in firing rates also follow a log-normal distribution. However, neurons are non-linear units of computation that utilize a wealth of nonlinear synaptic, somatic and dendritic mechanisms to transform synaptic input into output firing^14^, so weather changes follow a log-normal distribution is an empirical question which we should check. As expected, we find that also *changes* in firing rates during touch are log-normally distributed (see also Fig. S3a). If we plot our three metrics for changes in firing rate during social touch in somatosensory cortex, for example, we can see that the distributions of 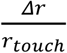 and 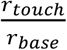 are skewed and not bell-shaped (they have a high skewness and a Gaussian distribution is a poor fit with a high negative log-likelihood). In contrast, the distribution of *β*_*touch*_ is approximately normal (the distribution has a low skewness and Gaussian distribution is a good fit, with a much lower negative log-likelihood):

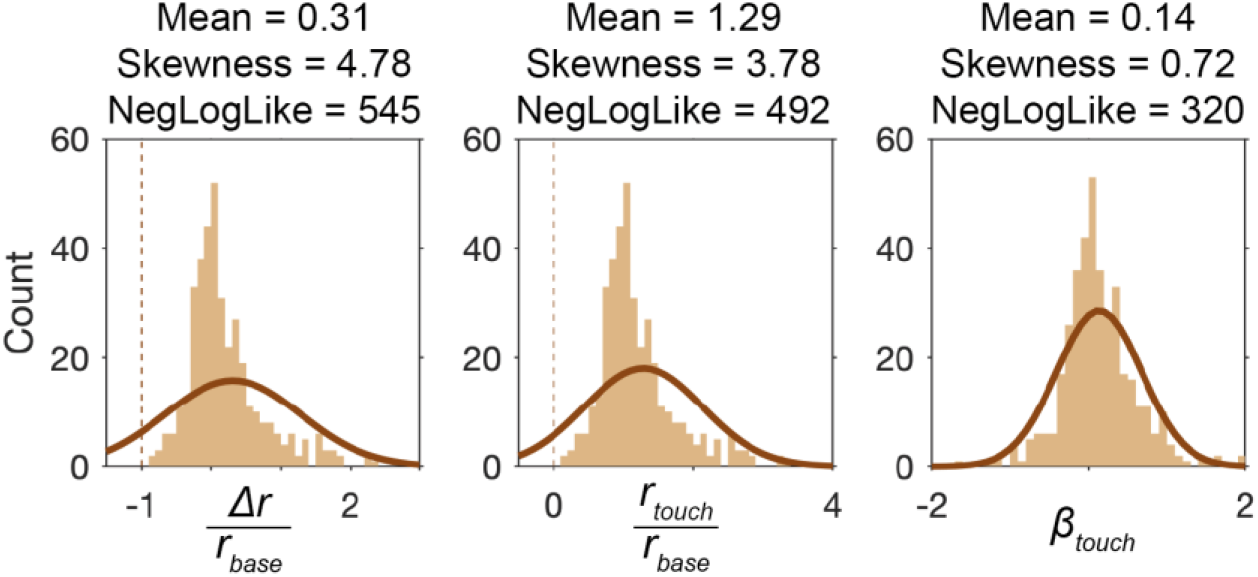

Ok, so both firing rates and *changes* in firing rates are approximately log-normally distributed. This is nice, since if we talk about an average *β*_*touch*_, for example, we are talking about a meaningful quantity, the mean of a bell-shaped distribution. The observant reader will note, however, that for generalized linear modeling, it’s not actually the variables, but rather the residuals of the variables that have to be normally distributed. So how do the residuals look? In our modeling, we want to model how the change in firing rates when touching female partners and touching male partners differs. If we plot both 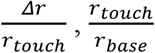 and *β*_*touch*_ for male and female partners (as in Fig 4. c-d) and fit a linear regression (top row below), they all suggest the same direction of the effect. However, when we inspect the residuals, we find that the regression residuals for 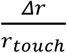 and 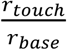 are quite skewed (Skewness > 1, meaning that assumptions of the the statistical model are violated), whereas the regression residuals for *β*_*beta*_. are approximately normal (meaning that the statistical model is valid).

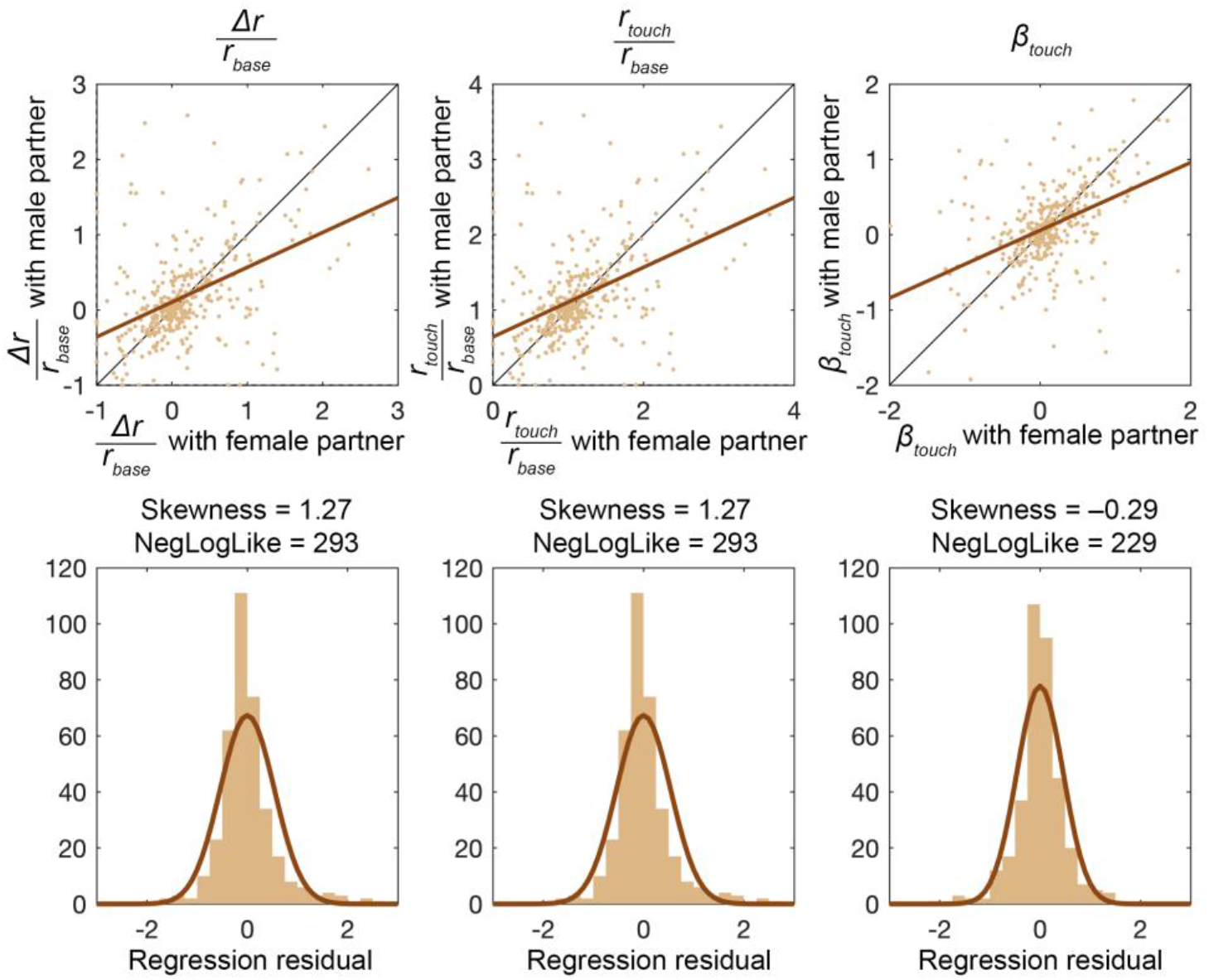

In summary, since both firing rates and changes in firing rates are log-normally distributed across cortical populations, *β*_*touch*_ is a natural metric for the network (the distribution is normal, so the mean and standard deviation is meaningful). Moreover, since the residuals of *β*_*touch*_ are approximately normal, statistical models relying on normally distributed errors are valid and we can use statistical modeling to quantify how the network firing patterns change with both social touch and social context.

### What is the connection to the regression model?

In our spike train regression, we model the firing rate of single neurons as a depending on spike history, recording session (to allow for baseline drift), touch and partner sex (see Methods). We assume that these effects are independent of each other, so we express the influence of the covariates on the instantaneous firing rate, *r*, by expressing the firing rate as a product of functions of the covariates^5^:

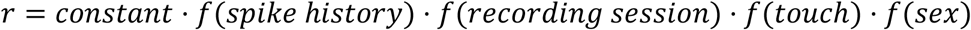

We are free to choose any functions *f*, as long as *f* ≥ 0 (firing rates cannot be negative) and *f* is injective (every *f(x)* is unique). The exponential function is a natural and convenient choice (*e*^*x*^ ≥ 0 for all *x* and *e*^*x*^ is injective). Moreover, exponential functions are convenient for computational reasons, since to calculate products of exponential functions, we just have to sum their exponents. In our model, touch and sex are binary variables, which we code as 0 and 1. In order to scale the relative impact of the various covariates, we need scaling factors, which we can pack inside the exponential function and call “*β*”. For example, if we disregard spike history, recording session and sex for now, we could write:

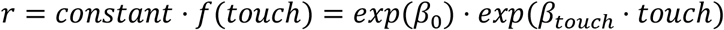

Touch is coded as a binary variable (0 or 1) so after we have used likelihood maximization to estimate the *β*-coefficients, we can calculate the baseline firing rate and the firing during touch as:

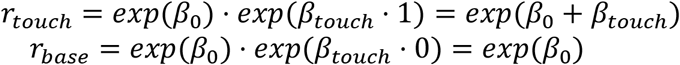

From this formulation, we can see that *β*_*touch*_ is the same as we introduced above:

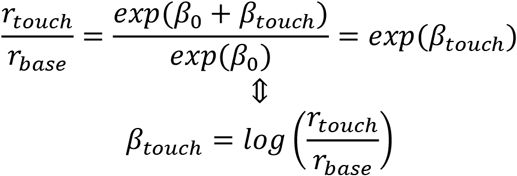

### Calculating responses with male and female interaction partners

In our modeling, we code periods of social touch as 0 and 1. If we do not model the effect of sex, then we can calculate the maximum likelihood estimate of the baseline and touch as shown above. In the models where we also fit the effect of sex, we code the data with an indicator variable such that *sex* = 1 corresponds to touching as male and *sex* = 0 corresponds to touching a female. In that way, we can fit a model where touch episodes with male and female conspecifics lead to different firing rate changes. If we again disregard spike history effects and baseline drift between recordings, we can calculate the firing rate when touching male and female conspecifics like this:

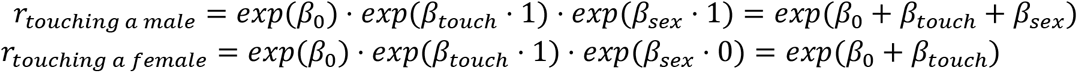

For example, if *β*_0_ = 0.7, *β*_*touch*_ = 0.1 and *β*_*sex*_ = 1.0, the neuron is slightly increased during social touch with females and strongly increased during social touch with males:

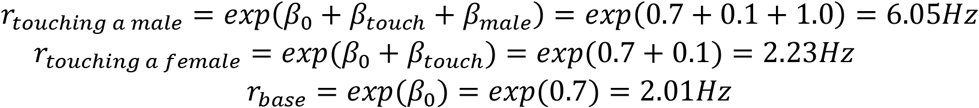

As another example, if *β*_0_ = 1.3, *β*_*touch*_ = 1.2 and *β*_*sex*_ = −1.2, the neuron is strongly increased during social touch with females and not modulated at all during social touch with males:

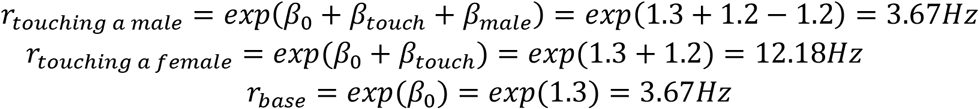

## Supplementary Note 2: Full specification of statistical models plotted in Fig. 4d & Fig. S5b

Full data and Matlab code required to fit and plot models are provided as supplementary data.

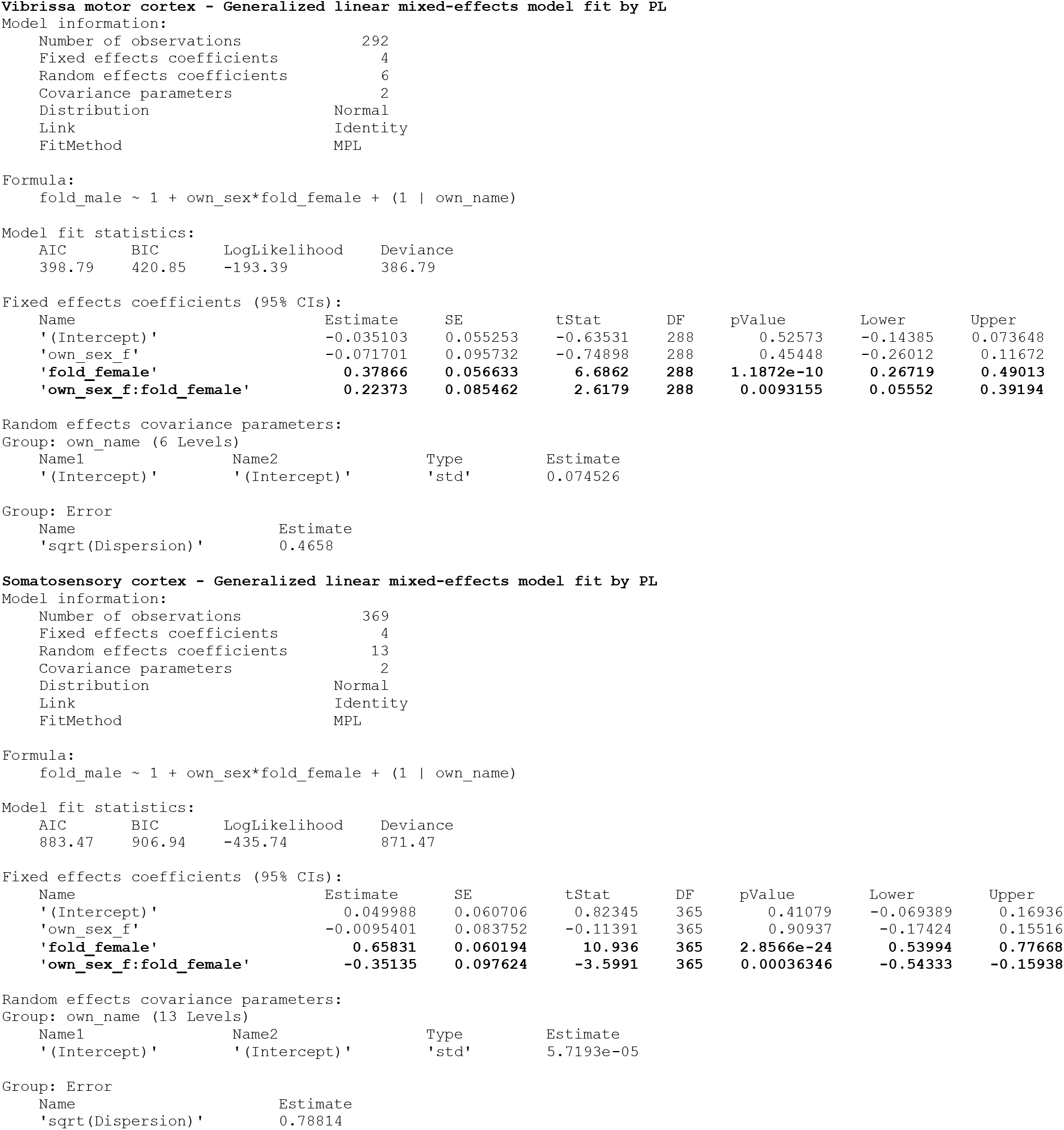
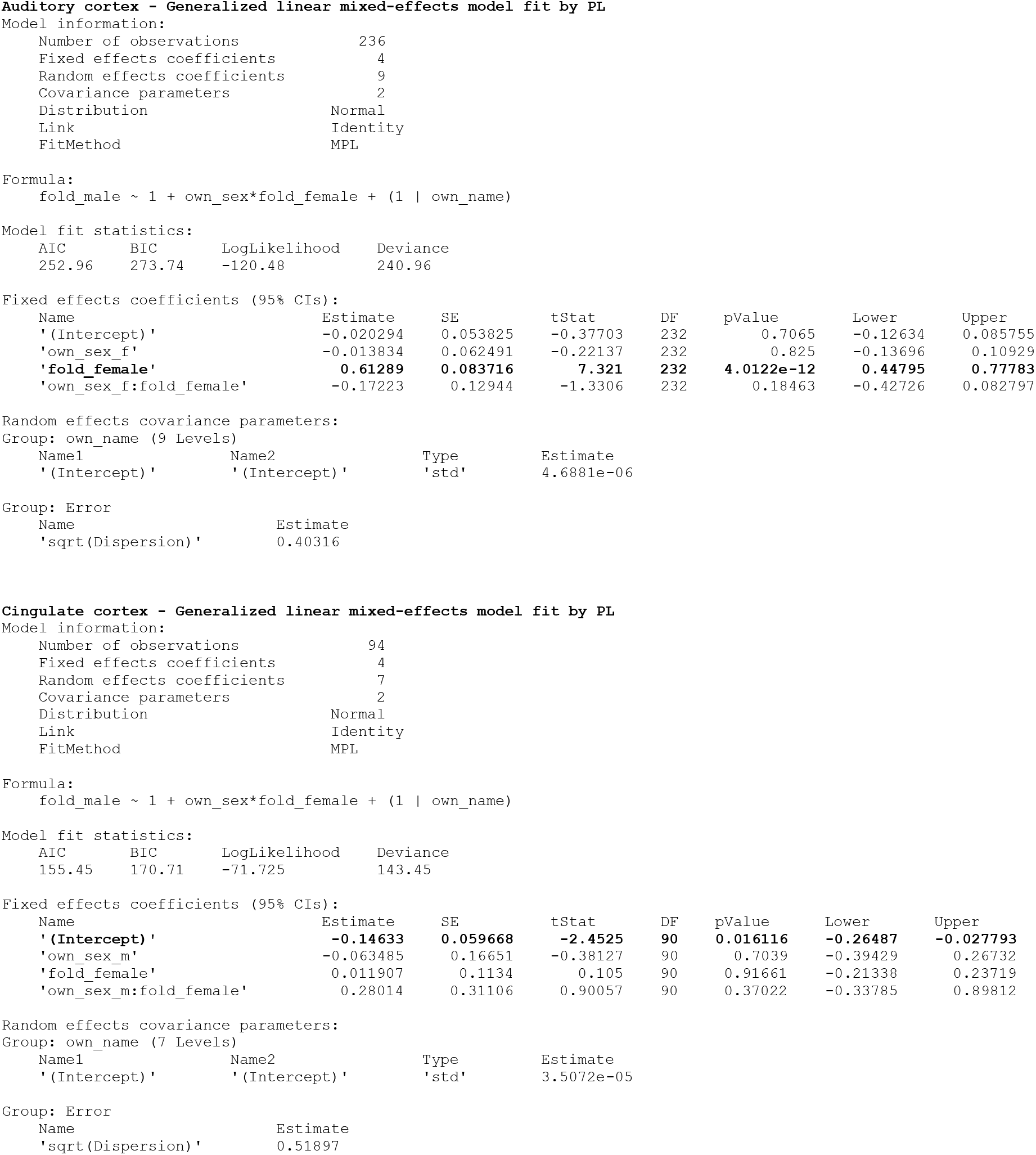
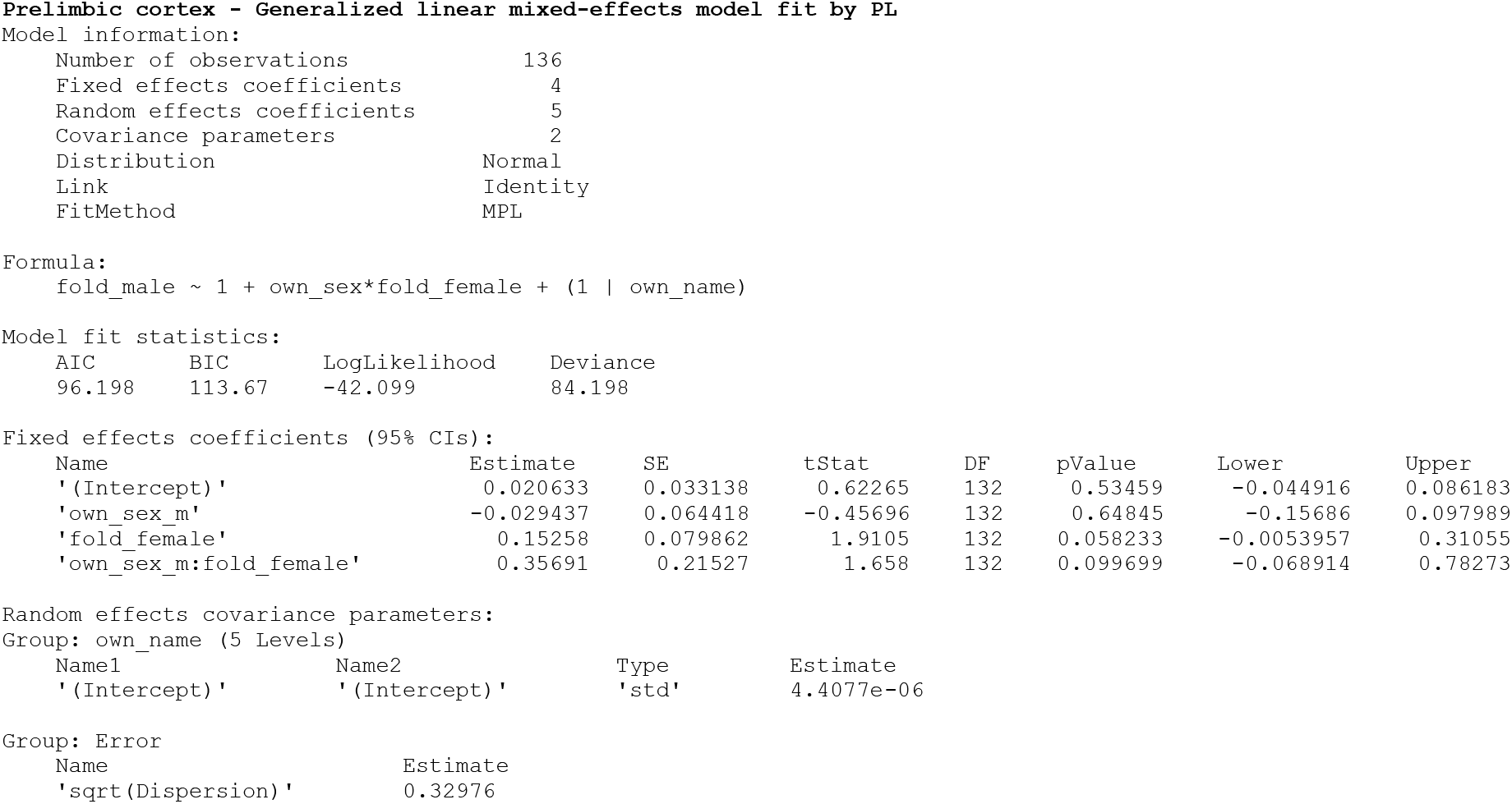

## Footnotes

1 Strictly speaking, if either the baseline or the touch rate is 0 Hz, log(*a*) is not defined, but tending to positive and negative infinity, respectively. This matches our intuition since, for example, total cessation of spiking is an “infinitely strong” decrease.

